# Fronto-central P3a to distracting sounds: an index of their arousing properties

**DOI:** 10.1101/333419

**Authors:** Rémy Masson, Aurélie Bidet-Caulet

## Abstract

The P3a observed after novel events is an event-related potential comprising an early fronto-central phase and a late fronto-parietal phase. It has classically been considered to reflect the attention processing of distracting stimuli. However, novel sounds can lead to behavioral facilitation as much as behavioral distraction. This illustrates the duality of the orienting response which includes both an attentional and an arousal component. Using a paradigm with visual or auditory targets to detect and irrelevant unexpected distracting sounds to ignore, we showed that the facilitation effect by distracting sounds is independent of the target modality and endures more than 1500 ms. These results confirm that the behavioral facilitation observed after distracting sounds is related to an increase in unspecific phasic arousal on top of the attentional capture. Moreover, the amplitude of the early phase of the P3a to distracting sounds positively correlated with subjective arousal ratings, contrary to other event-related potentials. We propose that the fronto-central early phase of the P3a would index the arousing properties of distracting sounds and would be linked to the arousal component of the orienting response. Finally, we discuss the relevance of the P3a as a marker of distraction.

## 1. Introduction

Attentional processes enable us to selectively attend stimuli which are relevant to our goals, and to filter out irrelevant stimuli, to increase task-efficiency. However, unexpected salient stimuli tend to attract our attention away from the task at hand: this is commonly referred as distraction or involuntary attention. This distraction effect is usually transient and we are able to focus back on the task, unless the stimulus was evaluated as significant.

Auditory distraction has been mostly investigated using audio-visual oddball paradigms during which task-irrelevant standard or rare novel sounds precede visual targets to be discriminated (Escera et al., 2003, 2000, 1998; Schröger and Wolff, 1998). These studies have shown that novel sounds, compared to standard sounds lead to prolonged reaction times or decreased hit rates to visual (Andrés et al., 2006; Berti et al., 2004; Escera et al., 1998) or auditory (Berti and Schröger, 2003; Rinne et al., 2006; Schröger, 1996; Wetzel et al., 2006) targets. These novel sounds would elicit attentional capture, also referred as involuntary orienting of attention in the literature (Wetzel et al., 2013), and their processing would require resources that are then unavailable for the maintenance of attention on the task at hand (Escera et al., 2000; Näätänen, 1992).

At the electrophysiological level, novel distracting sounds elicit a well-described sequence of event-related potentials (ERPs) which includes the N1 and a P3 complex (Escera et al., 2003, 1998). The N1 response to unexpected sounds is deemed to index a transient detector mechanism (Berti, 2013; Escera et al., 1998) that would trigger an attention switch to salient stimuli (Näätänen, 1990; Näätänen and Picton, 1987; Näätänen and Winkler, 1999). The P3 complex, also called novelty-P3 or P3a (Friedman et al., 2001; Olofsson and Polich, 2007; Ranganath and Rainer, 2003), is composed of a fronto-central early phase peaking around 235 ms, referred as the early-P3 in the following, and a fronto-parietal late phase peaking around 320 ms referred as the late-P3 in the following (Escera et al., 2000, 2000; Escera and Corral, 2007; Yago et al., 2003). This dissociation has been recently confirmed using a Principal Component Analysis (PCA) compiling data from different oddball paradigms (Barry et al., 2016). The functions attributed to each phase are still under debate. Nevertheless, the P3a has been considered to reflect the attention processing of distracting stimuli (Polich and Criado, 2006) and has been frequently associated with impaired performances due to unexpected novel sounds (Berti and Schröger, 2003; Escera et al., 1998; Schröger and Wolff, 1998; Wetzel et al., 2006), leading to the assumption that this ERP would index distraction.

However, in the last decades, a growing number of studies challenged the idea that unexpected rare novel sounds only yield distraction (for a review, see Parmentier, 2014). Indeed, a behavioral cost was not always observed after novels sounds (Li et al., 2013; Ljungberg et al., 2012; Parmentier et al., 2011, 2010; Wetzel et al., 2013, 2012). In some oddball paradigms, novel sounds could even enhance performances (SanMiguel et al., 2010b, 2010a; Wetzel et al., 2013, 2012). Several mechanisms can account for such facilitation effects in the oddball paradigms. Novel sounds could act as warning signals which trigger a top-down anticipation of the onset of the incoming target. They could also induce a phasic increase in unspecific arousal (SanMiguel et al., 2010b, 2010a; Wetzel et al., 2013, 2012), which results in enhanced readiness to process and respond to any upcoming stimulus. These findings led to propose that novel sounds generate a combination of facilitation and distraction effects which final effect on the performance of an unrelated task depends on the level of the task demands (SanMiguel et al., 2010a, see also Eysenck, 2012; Kahneman, 1973; Yerkes and Dodson, 1908 for the interaction of arousal with task demands) and the properties of the novel sounds (Parmentier et al., 2010; SanMiguel et al., 2010a; Wetzel et al., 2012).

The “arousal hypothesis” is consistent with a model of the orienting response towards unexpected novel sounds which comprises an arousal component to account for behavioral benefits and an attentional component, a reflexive attentional switch to the eliciting stimulus, to explain behavioral costs (Näätänen, 1992). Nevertheless, only few studies have investigated the link between the behavioral benefit and the properties of the distracting sounds (Max et al., 2015; Wetzel et al., 2012). To our knowledge, the differential impact of arousal and emotional properties on the facilitation effect has not been explored. We define here arousal as a state of enhanced physiological reactivity that leads to a condition of unspecific sensory alertness and readiness to respond to any upcoming stimulus (Aston-Jones and Cohen, 2005; Coull, 1998; Näätänen, 1992; Nieuwenhuis et al., 2011; Sturm and Willmes, 2001), enabling improved performance, in this particular case target detection.

In the same line, the P3 complex has been observed in response to novel sounds resulting in facilitation effects (SanMiguel et al., 2010b; Wetzel et al., 2013), leading to alternative explanations of the cognitive functions underlined by this brain response. Wetzel and colleagues proposed that the P3 complex reflects novelty evaluation of the stimulus; whereas San Miguel and colleagues proposed that the P3 complex reflects the summation of different brain operations triggered by the novel event, such as arousing, orienting and executive control processes. Accordingly, if we consider the different P3a phases, the early-P3 has not been necessarily linked to distraction in the literature. In oddball paradigms, in contrast to the late-P3, the early-P3 amplitude remains unchanged when behavioral distraction increases (SanMiguel et al., 2008), leading the authors to propose that the early-P3 may reflect a process other than the involuntary orienting of attention. In a similar fashion, behavioral distraction by deviant sounds is not always associated with the elicitation of an early-P3 (Horváth et al., 2011). On the other hand, the late-P3 has been more consistently considered as a signature of involuntary attentional capture (Bidet-Caulet et al., 2015; Escera et al., 2000, 1998; Roye et al., 2007). This undermines the idea that the P3a indexes solely distraction. It is imaginable that other properties of novel or distracting sounds (such as their arousing value) are linked to the P3a and especially to its early phase, the early-P3.

The goal of this study is to characterize the facilitation effect triggered by unexpected salient stimuli at the behavioral and electrophysiological levels. The Bidet-Caulet et al. (2015) paradigm is particularly adapted to investigate the duality of effects by distracting sounds. In this task, unexpected task-irrelevant sounds, played between a visual cue and an auditory target in 25% of the trials (see Fig. 1a), resulted in two opposite effects, whose intensity was found to be dependent on the delay between the distractor and the target (see Fig. 2). Distracting sounds played long before the target induce a reduction of reaction times compared to a condition without distractor: this difference has been considered as a behavioral index of facilitation (Bidet-Caulet et al., 2015). Distracting sounds played just before the target induce an increase in reaction times compared to a condition with a distractor played earlier: this difference has been considered as a behavioral index of attentional capture (Bidet-Caulet et al., 2015). Moreover, this paradigm produces a robust sequence of event-related potentials including a biphasic P3a.

**Fig. 1.**
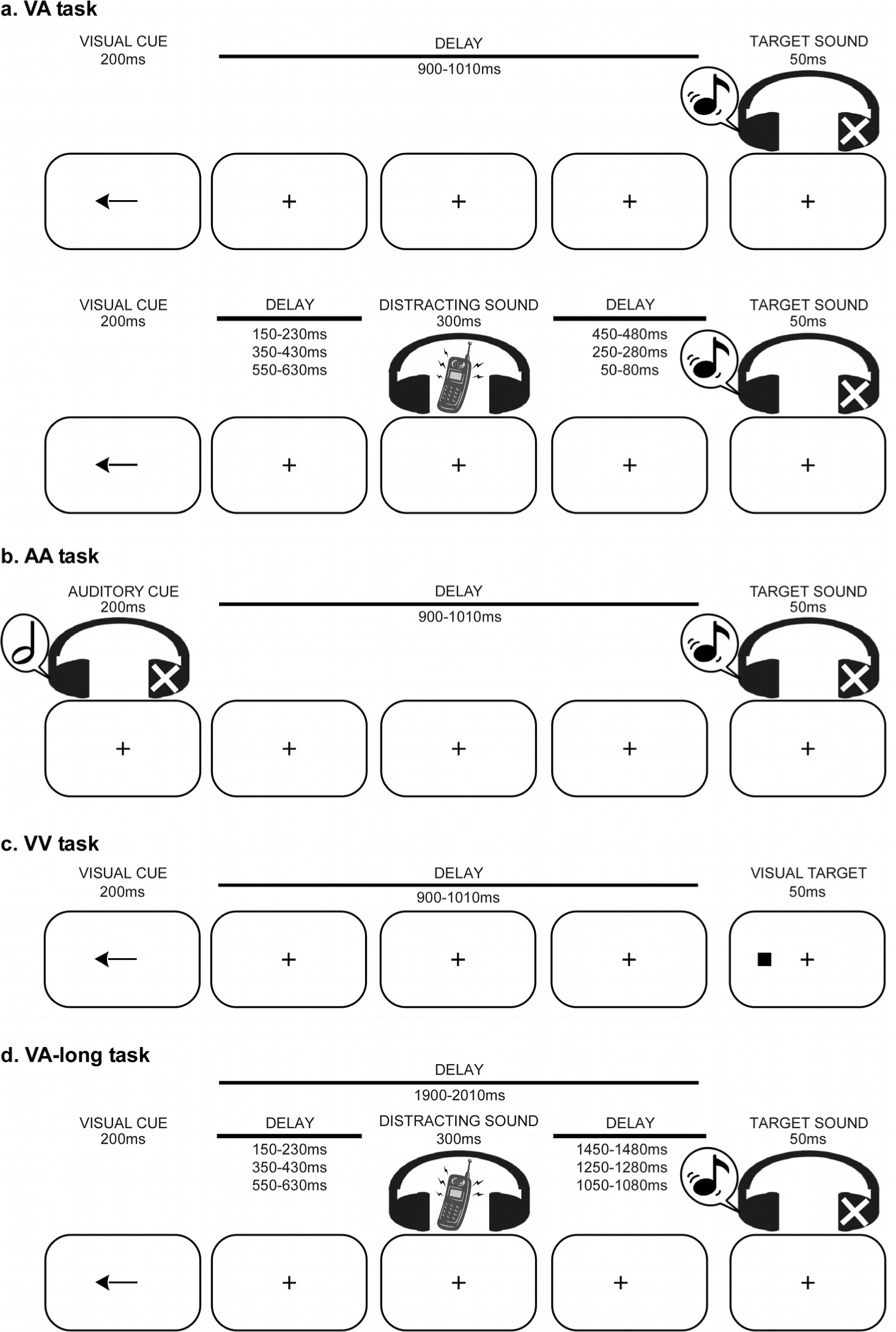
Experimental paradigms. **a. VA task.** First row: Example of a distractor-free trial (75% of the trials). A visual cue (200 ms) precedes the onset of a monaural target sound (50 ms) by a random delay (900-1010 ms). In 66.6% of the trials, a one-sided cue indicates in which ear (left or right) the target sound will be played (informative trial). In the other 33.4 % of the trials, a two-sided visual cue does not provide any indication in which ear (left or right) the target sound will be played (trial not depicted in the figure). Second row: Example of a trial with a distractor (25% of the trials). All parameters are identical to trials with no distracting sound, except that a binaural distracting sound (300 ms duration), such as a phone ring, is played during the delay between cue and target. The distracting sound can equiprobably onset in three different time periods after the cue offset: in the 150-230 ms range, in the 350-430 ms range, or in the 550-630 ms range. **b. AA task.** Example of a distractor-free trial. All parameters are identical to the VA task, except that the cue is a monaural (informative cue) or binaural (uninformative cue) 200 ms sound. **c. VV task.** Example of a distractor-free trial. All parameters are identical to the VA task, except that the target is a squared dot presented at the left or the right side of the screen. **d. VA-long task.** Example of a trial with a distractor. All parameters are identical to the VA task, except that the cue-target delay is comprised between 1900 and 2010 ms. VA: Visuo-auditory, AA: Audio-auditory, VV: Visuo-visual.

**Fig. 2.**
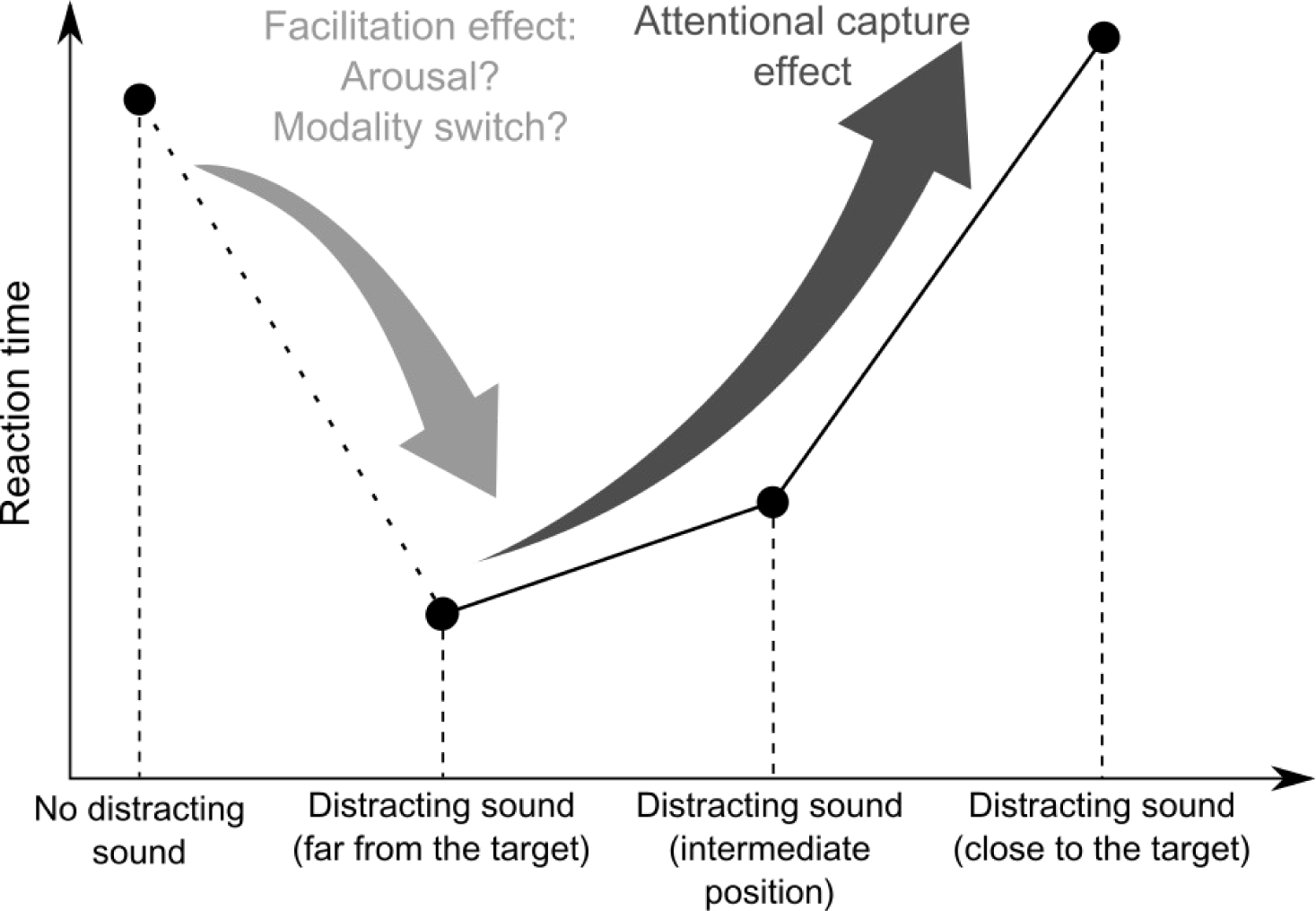
Schematic representation of the effect of distracting sounds on reaction times in a target detection task according to Bidet-Caulet et al. (2015). In comparison with trials without distractor, when the distractor-target delay is large, the presence of a distracting sound results in shorter reaction times (facilitation effect). However, when the distractor-target delay is short, this enhancement of performances by the distracting sound is canceled out (attentional capture effect).

Two hypotheses can account for the facilitation effect observed using this paradigm. As mentioned above, distracting sounds could trigger an increase in unspecific arousal. Alternatively, the auditory distractors could result in an earlier attentional shift from the visual cue modality to the auditory target modality, facilitating the processing of the target. In a first behavioral experiment, we used adaptations of the paradigm to test these two propositions by changing the sensory modality of the cue or the target (see Fig. 1b & c). In a second behavioral experiment, we used another adaptation of the paradigm to characterize the durability of the facilitation effect by increasing the delay between the distractor and the target (see Fig. 1d). Finally, in an EEG experiment, we investigated the brain underpinnings of the facilitation effect by exploring the relationship between the arousing properties of distracting sounds and distractor-related ERPs.

## 2. First behavioral experiment

In a recent study, Bidet-Caulet and colleagues managed to temporally dissociate facilitation and distraction components elicited by unexpected task-irrelevant sounds (Bidet-Caulet et al., 2015). In this task, a visual cue indicated (or not) the side of an auditory target to detect. Between the cue and the target, a distracting sound was presented in 25% of the trials (Fig. 1a). Distracting sounds, played more than 500 ms before the target, induced a reduction in reaction times compared to a condition without distractor. Two hypotheses can account for this facilitation effect. (1) The facilitation effect is related to an unspecific burst of arousal triggered by the distracting sound because of its unexpectedness and its saliency. (2) Facilitation is due to the distracting sound inducing an earlier attentional shift from the cue modality (visual) to the target modality (auditory, same as the distractor).

In this first behavioral experiment, we created two adaptations of the Bidet-Caulet et al. (2015) paradigm to test these hypotheses (Fig. 1b & c). In a visuo-visual task, we used a visual target instead of an auditory target. In an audio-auditory task, we used an auditory cue instead of a visual cue. We hypothesized that (1) if the behavioral facilitation effect by distracting sounds is caused by an increase in phasic arousal, this effect should not be influenced by the sensory modality of the cue nor of the target. However, (2) if the behavioral facilitation effect stems from an earlier attentional shift towards the auditory modality following distracting sounds, it should be reduced with an auditory cue or a visual target.

### 2.1 Materials and Methods

#### 2.1.1 Participants

Fifteen paid participants (all right-handed, 9 females, mean age ± standard deviation (SD): 28.5 ± 8.9 years) took part to the first behavioral experiment. All participants were free from neurological or psychiatric disorder, and had normal hearing and normal or corrected-to-normal vision. All participants gave written informed consent.

#### 2.1.2 Design of distracting sounds

Thirty ringing sounds (clock-alarm, door bell, phone ring, etc.) were used as distracting sounds in this study. These sounds were from several origins: Anne Guillaume’s sound library (Guillaume et al., 2004); Michael Marcell’s collection of environmental sounds (Marcell et al., 2000); audio CDs: sound library sonoteca, Mon imagier sonore (Olivier Tallec, Gallimard Jeunesse), Sound effects (DOM); and various websites: http://www.findsounds.com/, http://www.sound-effects-library.com/, and http://www.sounddogs.com/. When necessary, sounds were re-sampled at 44 kHz on 16 bits and edited to last 300 ms (5ms rise-time and 20ms fall-time), with Adobe Audition software (Adobe). All sounds were then RMS normalized with MATLAB (Mathworks). Finally, loudness was subjectively equalized by two listeners. Sounds were delivered at an intensity level judged comfortable by the participants at the beginning of the experiment.

#### 2.1.3 Stimuli and task

We manipulated the sensory modality of the cue and the target in three different cueing tasks (Fig.1a,b,c). In the visuo-auditory task (VA, see Fig 1a), 75 % of the trials (NoDIS trials) consisted in a visual cue (200 ms duration), a delay (randomly chosen between 900 and 1,010 ms) followed by a target sound (50 ms duration). The cue was centrally presented on a screen (grey background) and could be a green arrow pointing to the left, to the right, or to both sides. The target sound was a monaural harmonic sound (fundamental frequency: 200 Hz, 5 harmonics; 5 ms rise-time, 5 ms fall-time) presented at 15 dB SL (mean ± SD: 52.2 ± 0.7 dBA) in earphones. In the other 25 % (DIS trials), the same trial structure was used, but a binaural distracting sound at 35 dB SL (mean ± SD: 77.2 ± 0.7 dBA) was played during the delay. The cue and target categories were manipulated in same proportion for trials with and without distracting sound. In 33.3 % of the trials, the cue was pointing left and the target sound was played in the left ear, and in 33.3 % of the trials, the cue was pointing right and the target sound was played in the right ear, leading to a total of 66.6 % of informative trials. In the last 33.4 % of the trials, the cue was uninformative, pointing in both directions, and target sound was played in the left (16.7 %) or right (16.7 %) ear. The distracting sound could be equiprobably presented in three different time periods after the cue offset: in the 150–230 ms range (DIS1), in the 350–430 ms range (DIS2), or in the 550–630 ms range (DIS3).

Participants were instructed to perform a detection task by pressing a mouse button as fast as possible when they heard the target sound. They were asked to allocate their attention to the cued side in the case of informative cue. Participants were informed that informative cues were 100 % predictive and that a distracting sound could be sometimes played. In the absence of the visual cue, a blue fixation cross was presented at the center of the screen. Participants were instructed to keep their eyes fixating on the cross and to minimize eye movements and blinks while performing the task.

Two other variants of the task were designed. The structure remained the same but the sensory modality of the cue or of the target was modified. In the audio-auditory task (AA, see Fig. 1b), the cue was a harmonic sound (200 ms duration, 25 dB SL (mean ± SD: 67.2 ± 0.7 dBA), fundamental frequency: 500 Hz, 5 harmonics; 5 ms rise-time, 5 ms fall-time) played at the left or at the right ear (informative trials) or in both ears (uninformative trials). In the visuo-visual task (VV, see Fig. 1c), the target was a gray squared dot presented at the left or at the right side of the screen (visual angle: 6°) displayed for 50 ms.

Effects of the informational value of the cue are beyond the scope of this article and will not be presented. For a discussion of top-down attention deployment during the VA task, see Bidet-Caulet et al. (2015).

#### 2.1.4 Procedure

Participants were seated in a comfortable armchair in a sound attenuated room, at a 1.5 m distance from the screen. All stimuli were delivered using Presentation software (Neurobehavioral Systems, Albany, CA, USA). Sounds were delivered through earphones (SONY MDR-E819V). First, the auditory threshold was determined for the target sound, in each ear, for each participant using the Bekesy tracking method. This resulted in an average target threshold across subjects of 28.2 dBA (SD=0.6). Second, participants were trained with a short sequence of the task for each task.

Participants performed 5 blocks (72 trials each) for each of the tasks. Participants were thus presented, in each task, with 270 NoDIS, 30 DIS1, 30 DIS2, 30 DIS3 trials, irrespective of the cue category. No EEG was recorded in this experiment. The order of the tasks was randomly chosen for each participant. Participants had 2,500 ms to answer after targets.

#### 2.1.5 Behavioral data

A button press before target onset was considered as a false alarm (FA). A trial with no button press after target onset and before the next cue onset was considered as a missed trial. A trial with no FA and with a button press after target onset was counted as a correct trial. Reaction-times (RTs) to targets were analyzed in the correct trials only. Based on previous results (Bidet-Caulet et al., 2015), the difference in RTs between the NoDIS and DIS1 conditions can be considered as a measure of the facilitation effect triggered by distracting sounds whereas the difference in RTs between the DIS3 and DIS1 conditions can be considered as a good approximation of the attentional capture effect.

#### 2.1.6 Statistical Analysis of Behavioral data

Statistical analyses were conducted using the software JASP (Wagenmakers et al., 2018). RTs and percentages of correct trials were submitted to a Bayesian repeated-measure ANOVA with DISTRACTOR (four levels: NoDIS, DIS1, DIS2, DIS3) and MODALITY (three levels: AA, VA, VV) as within-participant factors. Post-hoc comparisons (based on the default t-test with a Cauchy prior) between levels of the MODALITY and of the DISTRACTOR factors were also conducted using the software JASP.

Effects of MODALITY on the facilitation effect (RT_NoDIS_ – RT_DIS1_) and the attentional capture effect (RT_DIS3_ – RT_DIS1_) were investigated as planned post-hoc analyses of the MODALITY by DISTRACTOR interaction, using Bayesian one-way ANOVA with MODALITY (three levels: AA, VA, VV) as the within-participant factor. Bayesian t-tests were also used to investigate the difference between the attentional capture effect and zero in each task.

We reported Bayes Factor (BF10) as a relative measure of evidence. To interpret the strength of evidence **against the null model**, we considered a BF between 1 and 3 as weak evidence, a BF between 3 and 10 as positive evidence, a BF between 10 and 100 as strong evidence and a BF higher than 100 as a decisive evidence (Lee and Wagenmakers, 2014). Similarly, to interpret the strength of evidence **in favor of the null model**, we considered a BF between 0.33 and 1 as weak evidence, a BF between 0.1 and 0.33 as positive evidence, a BF between 0.001 and 0.01 as strong evidence and a BF lower than 0.001 as a decisive evidence. The BF_inclusion_ compares models that contain the effect to equivalent models stripped of the effect and was considered as a relative measure of evidence supporting the inclusion of a factor. Detailed output of the Bayesian ANOVA are reported in the Table 1. Unless stated otherwise, in the “Results” section, mean values and standard errors of the mean are indicated.

**Table 1:** Results of Bayesian repeated-measure ANOVA on correct responses percentages, reaction times, facilitation effect and capture effect in the three experiments. The following outputs are specified: P(M)=prior model probability, P(M|data)=posterior model probability, BF_M_=change from prior to posterior model, BF_10_=Bayes Factor against the null model. The null model includes the subjects factor. For two-way Bayesian repeated-measure ANOVA, the following outputs are also specified: P(incl)=prior inclusion probability, P(incl|data)=posterior inclusion probability, BF_inclusion_=compares models that contain the effect to equivalent models stripped of the effect.

First behavioral experiment
| Models (Correct responses) | P(M) | P(M\|data) | BF<sub>M</sub> | BF<sub>10</sub> |
| --- | --- | --- | --- | --- |
| Null model | 0.20 | 0.11 | 0.48 | 1.00 |
| MODALITY | 0.20 | 0.02 | 0.08 | 1.05 |
| DIS | 0.20 | 0.67 | 8.00 | 6.10 |
| MODALITY + DIS | 0.20 | 0.14 | 0.66 | 2.21 |
| MODALITY + DIS + MODALITY * DIS | 0.20 | 0.06 | 0.27 | 0.88 |

| Models (Reaction times) | P(M) | P(M\|data) | BF<sub>M</sub> | BF<sub>10</sub> |
| --- | --- | --- | --- | --- |
| Null model | 0.20 | 1.77E-16 | 7.08E-16 | 1.00 |
| MODALITY | 0.20 | 3.65E-13 | 1.46E-12 | 2062.80 |
| DIS | 0.20 | 1.53E-06 | 6.12E-06 | 8.65E+09 |
| MODALITY + DIS | 0.20 | 0.31 | 1.76 | 1.73E+15 |
| MODALITY + DIS + MODALITY * DIS | 0.20 | 0.69 | 9.07 | 3.92E+15 |

| Models (Facilitation) | P(M) | P(M\|data) | BF<sub>M</sub> | BF<sub>10</sub> |
| --- | --- | --- | --- | --- |
| Null model | 0.50 | 0.38 | 0.62 | 1.00 |
| MODALITY | 0.50 | 0.62 | 1.62 | 1.62 |

| Models (Capture) | P(M) | P(M\|data) | BF<sub>M</sub> | BF<sub>10</sub> |
| --- | --- | --- | --- | --- |
| Null model | 0.50 | 0.05 | 0.05 | 1.00 |
| MODALITY | 0.50 | 0.95 | 19.71 | 19.71 |

| Effects (Correct responses) | P(incl) | P(incl\|data) | BF<sub>inclusion</sub> |
| --- | --- | --- | --- |
| MODALITY | 0.40 | 0.16 | 0.21 |
| DIS | 0.40 | 0.81 | 6.29 |
| MODALITY * DIS | 0.20 | 0.06 | 0.44 |

| Effects (Reaction times) | P(incl) | P(incl\|data) | BF<sub>inclusion</sub> |
| --- | --- | --- | --- |
| MODALITY | 0.40 | 0.31 | 200057.16 |
| DIS | 0.40 | 0.31 | 8.38E+11 |
| MODALITY * DIS | 0.20 | 0.69 | 2.27 |

Second behavioral experiment
| Models (Correct responses) | P(M) | P(M\|data) | BF<sub>M</sub> | BF<sub>10</sub> |
| --- | --- | --- | --- | --- |
| Null model | 0.20 | 0.39 | 2.54 | 1.00 |
| DELAY | 0.20 | 0.28 | 1.53 | 0.71 |
| DIS | 0.20 | 0.18 | 0.87 | 0.46 |
| DELAY + DIS | 0.20 | 0.13 | 0.61 | 0.34 |
| DELAY + DIS + DELAY * DIS | 0.20 | 0.02 | 0.10 | 0.06 |

| Models (Reaction times) | P(M) | P(M\|data) | BF<sub>M</sub> | BF<sub>10</sub> |
| --- | --- | --- | --- | --- |
| Null model | 0.20 | 1.66E-05 | 6.64E-05 | 1.00 |
| DELAY | 0.20 | 1.29E-05 | 5.14E-05 | 0.77 |
| DIS | 0.20 | 0.08 | 0.36 | 4965.79 |
| DELAY + DIS | 0.20 | 0.11 | 0.47 | 6356.66 |
| DELAY + DIS + DELAY * DIS | 0.20 | 0.81 | 17.29 | 48951.17 |

| Effects (Correct responses) | P(incl) | P(incl\|data) | BF<sub>inclusion</sub> |
| --- | --- | --- | --- |
| DELAY | 0.40 | 0.43 | 0.72 |
| DIS | 0.40 | 0.31 | 0.47 |
| DELAY * DIS | 0.20 | 0.02 | 0.18 |

| Effects (Reaction times) | P(incl) | P(incl\|data) | BF<sub>inclusion</sub> |
| --- | --- | --- | --- |
| DELAY | 0.40 | 0.11 | 1.28 |
| DIS | 0.40 | 0.19 | 6379.94 |
| DELAY * DIS | 0.20 | 0.81 | 7.70 |

EEG experiment
| Models (Correct responses) | P(M) | P(M\|data) | BF<sub>M</sub> | BF<sub>10</sub> |
| --- | --- | --- | --- | --- |
| Null model | 0.20 | 1.70E-17 | 6.82E-17 | 1.00 |
| AROUSAL | 0.20 | 2.10E-18 | 8.41E-18 | 0.12 |
| DIS | 0.20 | 0.86 | 23.70 | 5.02E+16 |
| AROUSAL + DIS | 0.20 | 0.11 | 0.47 | 6.22E+15 |
| AROUSAL + DIS + AROUSAL * DIS | 0.20 | 0.04 | 0.16 | 2.25E+15 |

| Models (Reaction times) | P(M) | P(M\|data) | BF<sub>M</sub> | BF<sub>10</sub> |
| --- | --- | --- | --- | --- |
| Null model | 0.20 | 1.35E-49 | 5.41E-49 | 1.00 |
| AROUSAL | 0.20 | 1.91E-50 | 7.63E-50 | 0.14 |
| DIS | 0.20 | 0.85 | 23.38 | 6.31E+48 |
| AROUSAL + DIS | 0.20 | 0.14 | 0.65 | 1.04E+48 |
| AROUSAL + DIS + AROUSAL * DIS | 0.20 | 0.01 | 0.02 | 4.03E+46 |

| Effects (Correct responses) | P(incl) | P(incl\|data) | BF<sub>inclusion</sub> |
| --- | --- | --- | --- |
| AROUSAL | 0.40 | 0.11 | 0.13 |
| DIS | 0.40 | 0.96 | 4.99E16 |
| AROUSAL * DIS | 0.20 | 0.04 | 0.36 |

**Table 1:** Results of Bayesian repeated-measure ANOVA on correct responses percentages, reaction times, facilitation effect and capture effect in the three experiments.
| Effects (Reaction times) | P(incl) | P(incl\|data) | BF<sub>inclusion</sub> |
| --- | --- | --- | --- |
| AROUSAL | 0.40 | 0.14 | 0.16 |
| DIS | 0.40 | 0.99 | 6.44E+48 |
| AROUSAL * DIS | 0.20 | 5.46E-3 | 0.04 |

### 2.2 Results

#### 2.2.1 Percentage of correct responses (Table 1)

Participants correctly performed the detection task in 97.7% ± 1.1% of the trials. The remaining trials were either missed trials (0.1%) or trials with FAs (1.9% ± 0.3%).

For the percentage of correct trials, the only model showing some evidence only included the effect of DISTRACTOR (BF_10_=6.1). There was positive evidence against a specific effect of MODALITY (BF_inclusion_=0.2) and there was positive evidence of a specific effect of DISTRACTOR (BF_inclusion_=6.3) on the percentage of correct trials. Post-hoc comparisons showed that participants had a slightly lower percentage of correct responses in the DIS1 condition (95.3% ± 1.5%) compared to the NoDIS condition (98.1% ± 0.4%) (BF_10_=17.3).

#### 2.2.2 Reaction times (Fig. 3, Table 1)

The best model was the one with both main effects of DISTRACTOR and MODALITY and their interaction (BF_10_=3.9·10^15^), but it was only 2.3 times more likely than the model with only the two main effects of DISTRACTOR and MODALITY (BF_10_=1.7·10^15^). There was decisive evidence for specific effects of DISTRACTOR (BF_inclusion_=8.4·10^11^) and MODALITY (BF_inclusion_=2.0·10^5^), there was only weak evidence for a specific effect of the DISTRACTOR by MODALITY interaction (BF_inclusion_=2.3).

**Fig. 3.**
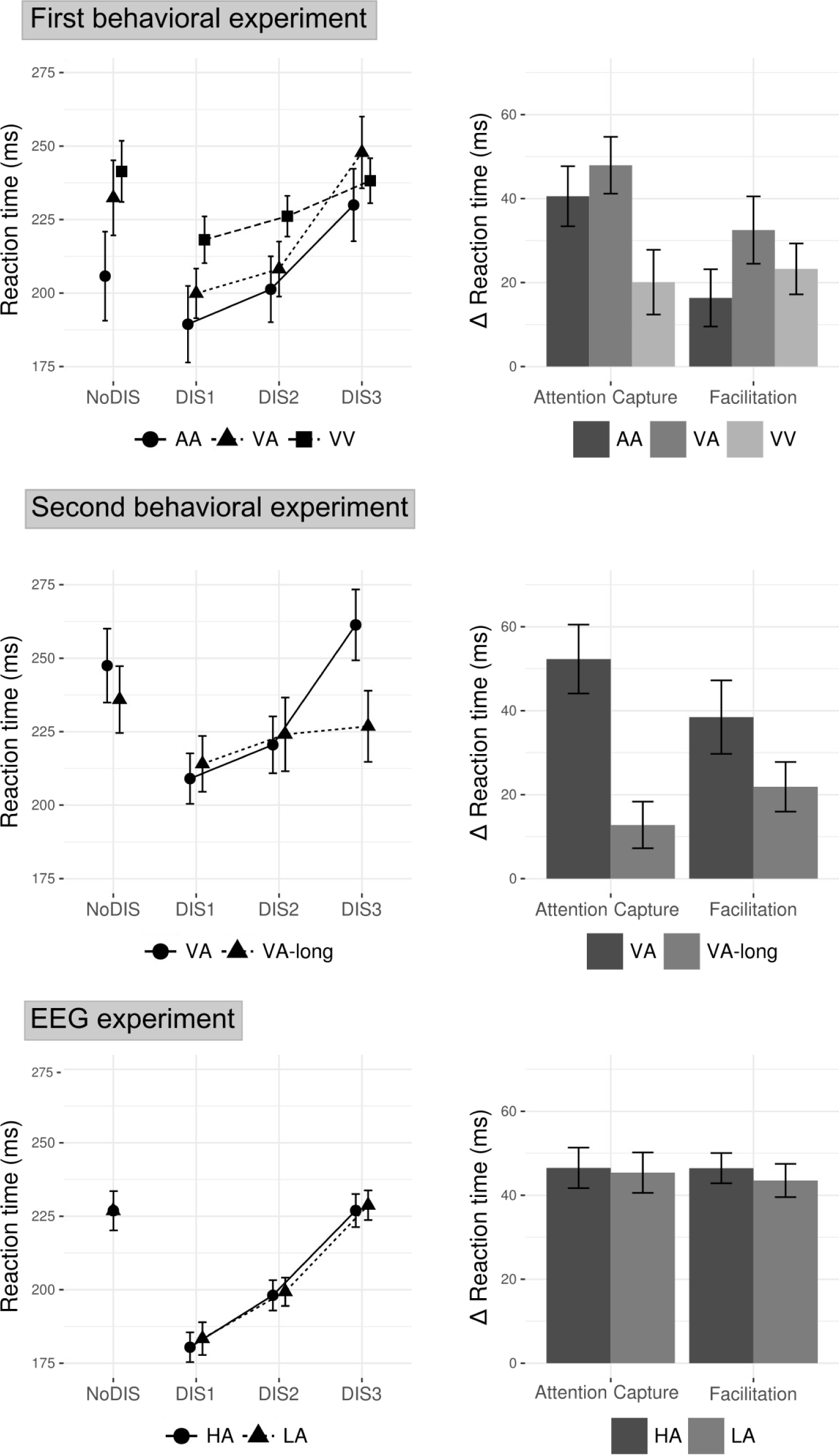
Behavioral results for the three experiments. **Left.** Mean reaction time (RT) as a function of the distractor condition (NoDIS, DIS1, DIS2, DIS3). **Right.** Attentional capture (RT to DIS3 minus RT to DIS1) and facilitation (RT to NoDIS minus RT to DIS1) effects. Error bars represent +/- 1 SEM. VA: Visuo-auditory, AA: Audio-auditory, VV: Visuo-visual, LA = low arousing sounds, HA = high arousing sounds.

Post-hoc analysis of the MODALITY main effect showed that participants were faster in the AA task than in the VA (BF_10_(AA vs. VA)=1381.8) and VV (BF_10_(AA vs. VV)=21834.9) tasks.

Post-hoc analysis of the DISTRACTOR main effect showed that participants were faster in the DIS1 condition than in the NoDIS (BF_10_=1.7·10^5^) and DIS3 (BF_10_=2.7·10^8^) conditions, confirming the significance of a facilitation effect and an attentional capture effect. On average, between NoDIS and DIS1, RTs dropped by 9.5% ± 2.7%.

Planned post-hoc analyses of the DISTRACTOR by MODALITY interaction were performed to test the effect of MODALITY on the facilitation and attentional capture effects. Almost no evidence of an effect of MODALITY on facilitation (BF_10_=1.6) was found. A strong evidence of an effect of MODALITY on attentional capture (BF_10_=19.7) was found, with post-hoc comparisons showing strong evidence of a reduced attentional capture in the VV task compared to the VA task (BF_10_=11.3). No evidence of difference in attentional capture between VA and AA (BF_10_=0.8) was found. However, there were positive to decisive evidence of an attentional capture effect in all the tasks (Bayesian t-tests comparing the effect to zero: BF_10_(AA)=1270.6, BF_10_(VA)=9130.6, BF_10_(VV)=3.8).

### 2.3 Discussion

This first experiment confirms that distracting sounds can result in a facilitation effect at the behavioral level, and shows that this benefit is independent of the cue and target sensory modalities.

#### 2.3.1 Facilitation and attentional capture effects of distracting sounds

With the present protocol, it has been possible to temporally dissociate facilitation and attentional capture effects of the same distracting sounds within the same task. Distracting sounds elicit a facilitation effect that improves performances especially with long distractor-target intervals (i.e. DIS1 and DIS2 conditions). However, the gain in RT is canceled out when the distractor onset is too close to the target sound (DIS3 condition), suggesting that distractors also trigger an attentional capture. This timing effect indicates that when the target occurs, the balance between cost and benefit differs according to the onset time of the distracting sound, suggesting that the facilitation and attentional capture effects triggered by the distracting sounds have different time courses. The facilitation effect would be stable for at least 750 ms (longest interval between distracting and target sound onsets) and induce a similar benefit on target processing irrespective of the distractor onset time. On the contrary, the attentional capture would be a transient powerful phenomenon that could interfere with target processing only when the distracting sound occurs within a few hundreds of ms before target onset, and that would be low or completed when the distractor is played long before target onset. A short duration of the attentional capture phenomenon is consistent with previous findings showing a detrimental effect of task-irrelevant deviant sound on target processing with onset-to-onset interval of 200 ms but not of 550 ms (Schröger, 1996). According to our results, the facilitation effect of novel sounds is less likely to be observed, at the behavioral level, in classical oddball studies because the typical sound-target delay is 300 ms, thus the attentional capture effect has not yet vanished and masks any potential facilitation effect.

These present results confirm that an unexpected salient sound triggers several phenomena that may produce opposite effects on the reaction time to a subsequent target: a cost related to a short-lasting attentional capture and the benefits of a more long-lasting facilitation effect. This duality in effects was previously observed in several recent studies (Ljungberg et al., 2012; Max et al., 2015; Parmentier et al., 2010; SanMiguel et al., 2010b, 2010a; Wetzel et al., 2013, 2012).

#### 2.3.2 Nature of the facilitation effect

With this first experiment, we attempted to precise the nature of the facilitation effect. In the visuo-auditory task, two hypotheses may account for the gain in RT in distractor trials. (1) The facilitation effect is related to an unspecific burst of arousal triggered by the distracting sound because of its unexpectedness and its saliency and because of the low arousal level of the task, as proposed in previous studies (Bidet-Caulet et al., 2015; SanMiguel et al., 2010b, 2010a; Wetzel et al., 2013, 2012). (2) Facilitation is due to the facilitation by the distracting sound of the attentional shift from the cue modality (visual) to the target modality (auditory, same as the distractor). A distracting sound placed between a visual cue and an auditory target could result in an earlier attentional shift leading to better performances compared to a no distracting sound condition. According to this second hypothesis, we should expect a smaller behavioral benefit if the cue and the target are in the same sensory modality. We tested this hypothesis in a visuo-visual and an audio-auditory cueing tasks and found no clear evidence of a reduction of the facilitation effect triggered by distracting sounds, in comparison to the visuo-auditory task. This is in line with previous results that showed that auditory warning signals can trigger facilitation in both visual and auditory detection tasks (Posner et al., 1976). Therefore, the attentional reorientation towards the target modality cannot account for the facilitation effect of the distracting sound. Moreover, similar facilitation effects of the distracting sounds were found regardless of the target sensory modality, suggesting that the facilitation effect is independent of the sensory modality. These results rather support the first hypothesis that facilitation effect stems from a distractor-related burst of arousal.

Outside of the literature on the oddball paradigm, it has also been known for decades that auditory warning signals, also referred to as “accessory stimuli”, played before or concurrent to the onset of the target visual stimulus facilitate the processing of such target stimulus (Hackley and Valle-Inclán, 1999, 1998; Posner et al., 1976; Valls-Solé et al., 1995). Interestingly, it has also been proposed that the speeding of reaction times by warning signals/accessory stimuli stems from an increase in phasic arousal (Hackley, 2009; Tona et al., 2016).

In conclusion and in agreement with previous studies (SanMiguel et al., 2010b; Wetzel et al., 2013), we show that performances after a distracting sound result from the combination of the benefits due to an increase in unspecific arousal and to the costs of attentional capture.

#### 2.3.3 Effects of cue and target sensory modality on reaction times and attentional capture effects

On the one hand, subjects were faster in the audio-auditory task than in the visuo-auditory one. It could be easily explained by the fact that auditory cues are more arousing than the visual ones inducing a general gain in reactivity. On the other hand, subjects were slower overall in the visuo-visual task simply because the task was probably more difficult.

There was positive to decisive evidence of an attentional capture effect regardless of the sensory modality of the cue and target. This result does not support the hypothesis that, in cross-modal tasks, distraction stems from the time penalty caused by shifting attention from one modality to the other (Parmentier, 2014; Parmentier et al., 2008). The attentional capture effect would be rather due to an involuntary capture of attention by the distracting event, diverting attention from the task at hand (Escera et al., 2000; Horváth et al., 2008; Näätänen, 1992).

We also found that the attentional capture effect is weaker in the visuo-visual task, but comparable in the visuo-auditory and audio-auditory tasks. In the visuo-visual task, the auditory modality is not relevant for the task: as a result, subjects may be more efficient at inhibiting sounds, and less sensitive to attentional capture by distracting sounds.

## 3. Second behavioral experiment

The aim of this second behavioral experiment is to test the endurance of the facilitation effect: a reduction or even an extinction of the facilitation effect could be expected with time. We created an adaptation of Bidet-Caulet et al. (2015) paradigm with twice longer cue-target intervals (in order to increase the delay between the distracting sound and the target by 1 s, see Fig. 1d).

### 3.1 Material and methods

#### 3.1.1 Participants

Twelve paid participants (all right-handed, 4 females, mean age ± SD: 26 ± 5.4 years) took part to the second behavioral experiment. Six participants participated to both behavioral experiments.

All participants were free from neurological or psychiatric disorder, and had normal hearing and normal or corrected-to-normal vision. All participants gave written informed consent.

#### 3.1.2 Stimuli and task

Participants performed the VA task and a variant of this task (VA-long, see Fig. 1d). The order of the tasks was randomized between subjects. The task structure remained the same however in the VA-long task, the cue-target delay was increased by 1000 ms (between 1900 and 2010 ms). Cue-distractor delays were unchanged: therefore, the distractor-target interval was increased by 1000 ms. The experimental procedure was identical than during the first experiment. The auditory threshold was determined for the target sound, in each ear, for each participant using the Bekesy tracking method. This resulted in an average target threshold of 28.2 dBA (SD=0.7), in an average distractor loudness of 77.2 dBA (SD=0.7) and in an average target loudness of 52.2 dBA (SD=0.7)

#### 3.1.3 Statistical Analysis of Behavioral data

RTs and percentages of correct trials were submitted to a Bayesian repeated-measure ANOVA with DISTRACTOR (four levels: NoDIS, DIS1, DIS2, DIS3) and DELAY (two levels: VA, VA-long) as within-participant factors. Post-hoc comparisons (based on the default t-test with a Cauchy prior) between levels of the DELAY factor were also conducted using the software JASP.

Effects of DELAY on the facilitation effect (RT_NoDIS_ – RT_DIS1_) and the attentional capture effect (RT_DIS3_ – RT_DIS1_) were investigated as planned post-hoc analyses of the DELAY by DISTRACTOR interaction, using Bayesian t-tests. Bayesian t-tests were also used to investigate the difference between the attentional capture effect and zero in each task. Detailed output of the Bayesian ANOVA are reported in the Table 1. Unless stated otherwise, in the “Results” section, mean values and standard errors of the mean are indicated.

### 3.2 Results

#### 3.2.1 Percentage of correct responses (Table 1)

Participants correctly performed the detection task in 98.3% ± 0.4% of the trials. The remaining trials were either missed trials (0.1%) or trials with FAs (1.6% ± 0.4%). No model showed any evidence of an effect on the percentage of correct trials (all BF_10_<1).

#### 3.2.2 Reaction times (Fig. 3, Table 1)

The best model was the one with both main effects of DISTRACTOR and DELAY and their interaction (BF_10_=48951.2). There was a decisive specific effect of DISTRACTOR (BF_inclusion_=6379.9) and a positive specific effect of the DISTRACTOR by DELAY interaction (BF_inclusion_=7.7). However, there was no evidence for a specific effect of DELAY (BF_inclusion_=1.3).

Post-hoc analysis of the DELAY main effect did not show any evidence for a difference in RTs between the VA and VA-long task (BF_10_=0.7).

Planned post-hoc analyses of the DISTRACTOR by DELAY interaction were performed to test the effect of DELAY on the facilitation and attentional capture effects. We observed only weak evidence of an effect of DELAY on facilitation (BF_10_=2.0) but we observed strong evidence of an effect of DELAY on attentional capture (BF_10_=17.1). Attentional capture was lower in the VA-long task (13.6 ± 5.9 ms) compared to the VA task (50.0 ± 7.4 ms). There was not much evidence that attentional capture is different from zero in the VA-long task (Bayesian t-test: BF_10_=1.9), whereas there was decisive evidence that the effect is different from zero in the VA task (Bayesian t-test: BF_10_=505.6).

### 3.3 Discussion

In this second experiment, we wanted to characterize the durability of the facilitation effect triggered by distracting sounds. In the longer version of the experiment (VA-long), participants still responded faster to targets preceded by a distracting sound despite a distractor-target delay reaching up to 1750 ms. Even if a trend could be observed in the data, there is no positive evidence that the facilitation effect decreased with a longer delay. This increase in phasic arousal seems to remain sustained at a steady level for at least a couple of seconds. A facilitation effect of salient task-irrelevant sounds has already been observed for sound-target delay superior to 1000 ms (as well as for very short delays under 100 ms) in several studies (e.g. Posner et al., 1976; Ulrich, 1996). In a paradigm using auditory targets preceded by standard or novel pictures, novel pictures were found to reduce reaction times but only for picture-target delay of 0 and 200 ms and not for 800 ms (Schomaker and Meeter, 2014). This could be explained by the fact that the novel stimuli in the visual modality are less arousing than those in the auditory modality. Visual accessory stimuli trigger less facilitation than auditory accessory stimuli in both visual and auditory detection tasks (Posner et al., 1976).

These results also confirm that attentional capture is a transient phenomenon: when the distractor-target delay is superior to 1000 ms, no clear evidence could be found of a difference in reaction times between trials including early distractors and trials including late distractors. As discussed previously, this is consistent with results using the auditory oddball paradigm: a behavioral effect of attentional capture by deviant sounds can only be observed for short delays between the deviant and the target sound (200 ms) but not for longer delays (550 ms) (Schröger, 1996).

## 4. EEG experiment

In this EEG experiment, we investigated the influence of the arousing content of the distracting sounds on their respective event-related potentials during the Bidet-Caulet et al. original task (Fig. 1a). We hypothesize that the increase of phasic arousal following distracting sounds is reflected at some point in the cortical processing of the sound. In that case, we expected that there is at least one event-related response whose amplitude would be modulated by the perceived “arousingness” of the sound. In this purpose, distracting sounds were rated according to their arousing content and ERPs to high and low arousing sounds were compared. To dissociate the effect of “arousingness” from the impact of emotional valence, multiple regression were performed on ERPs with arousal and valence as regressors.

### 4.1 Materials and Methods

#### 4.1.1 Participants

Forty-one paid participants took part to the EEG experiment. Two of them were excluded from further analyses because of noisy EEG data, one because of low behavioral performances and one because of an issue with the experimental setup during the study. Consequently, only thirty-seven participants (21 females, all right-handed, mean age ± SD: 22.5 ± 2.7 years) were included in further analyses. Data from 17 out of the 37 included participants have been presented in a previous publication (Bidet-Caulet et al., 2015).

No participant participated in both EEG and behavioral experiments. All participants were free from neurological or psychiatric disorder, and had normal hearing and normal or corrected-to-normal vision. All participants gave written informed consent.

#### 4.1.2 Stimuli, task and procedure

The task was identical to the VA task (Fig. 1a) in behavioral experiments and to the experiment in Bidet-Caulet et al. (2015). EEG was recorded while participants performed 15 blocks (72 trials each) of the VA task. Participants were thus presented, with 810 NoDIS and 270 DIS trials. The experimental procedure was identical than during the previous experiments. The auditory threshold was determined for the target sound, in each ear, for each participant using the Bekesy tracking method. This resulted in an average target threshold of 28.9 dBA (SD=2.2), in an average distractor loudness of 77.9 dBA (SD=2.2) and in an average target loudness of 47.9 dBA (SD=2.2).

#### 4.1.3 Behavioral ratings of distracting sounds

The same set of distracting sounds used in the behavioral experiments were used in the EEG experiment. Forty-one participants were asked to rate each sound for arousal (calm – exciting) and for valence (unhappy – happy) on a 9-point scale with the Self-Assessment Manikins (Bradley and Lang, 1994). Twenty of these participants were from the cohort of the EEG experiment and the twenty-one additional ones were only recruited for the sound rating. The thirty distracting sounds were presented in a different random order to each participant. After each sound, participants were successively presented with the arousal and the emotion scales. They had 5s to indicate their rating value by pressing the corresponding key on a keyboard for each scale.

High-arousing (HA) distracting sounds will comprise those rated above the median rating of all sounds, low-arousing (LA) distracting sounds will comprise those rated beneath the median rating of all sounds. Acoustic parameters (mean brightness, mean roughness, mean pitch and attack leap) were analyzed using the MATLAB MIRtoolbox 1.3.4 (Lartillot et al., 2008).

#### 4.1.4 Statistical Analysis of Behavioral data

RTs and percentages of correct trials were submitted to a Bayesian repeated-measure ANOVA with DISTRACTOR (four levels: NoDIS, DIS1, DIS2, DIS3) and AROUSAL (two levels: HA, LA) as within-participant factors. Post-hoc comparisons (based on the default t-test with a Cauchy prior) between levels of the DISTRACTOR factor were also conducted using the software JASP. Detailed output of the Bayesian ANOVA are reported in the Table 1. Unless stated otherwise, in the “Results” section, mean values and standard errors of the mean are indicated.

#### 4.1.5 EEG Recording

EEG was recorded from 32 active Ag/AgCl scalp electrodes mounted in an electrode-cap (actiCap, Brain Products, Gilching, Germany) following a sub-set of the extended International 10–10 System. Four additional electrodes were used for horizontal (external canthi locations) and vertical (left supraorbital and infraorbital ridge locations) EOG recording and two other electrodes were placed on earlobes. The reference electrode was placed on the tip of the nose and the ground electrode on the forehead. Data were amplified, filtered and sampled at 1,000 Hz (BrainAmp, Brain Products, Gilching, Germany). Data were re-referenced offline to the average potential of the two earlobe electrodes.

#### 4.1.6 EEG Data Analysis

EEG data were band-pass filtered (0.5–40 Hz). Prior to ERP analysis, eye-related activities were detected using independent component analysis (ICA) and were selectively removed via the inverse ICA transformation. Only 1 or 2 ICs were removed in each participant. Trials including false alarms or undetected target, and trials contaminated with excessive muscular activity were excluded from further analysis. For the purpose of the present study, only trials with distracting sounds were considered (see Bidet-Caulet et al., 2015 for analysis of the other trials). On average across participants, the number of considered trials for analysis was 119 ± 12 trials (mean ± SD) with HA and 119 ± 11 trials with LA distracting sounds.

ERPs were averaged locked to distractor onset and were baseline corrected to the mean amplitude of the -100 to 0 ms period before distractor onset. To analyze ERPs to distracting sound, for each distractor onset time-range, surrogate ERPs were created in the NoDIS trials and subtracted from the actual ERPs. The obtained ERPs were thus clear of cue-related activity.

ERP scalp topographies were computed using spherical spline interpolation (Perrin et al., 1989). ERPs were analyzed using the software package for electrophysiological analysis (ELAN Pack) developed at the Lyon Neuroscience Research Center (; Aguera et al., 2011).

#### 4.1.7 Statistical Analysis of ERP data

Statistical analyses were performed on a frontal (Fz), a central (Cz) and a parietal (Pz) electrode from the distractor onset to 350 ms after onset as previously done in Bidet-Caulet et al., 2015. Later responses were not investigated because the shortest duration between the distracting sound and the following target sound onset was 350 ms.

To explore the effect of the arousing content on the ERPs to distracting sound (HA vs. LA), permutation tests based on randomization (Edgington, 2014) were used. Each randomization consisted in (1) the random permutation of the values of the 37 pairs of conditions (corresponding to the 37 participants), (2) the calculation of the sum of squared sums of values in each condition, (equivalent to the test statistic F in designs with paired conditions with equal sample size, for a demonstration see Edgington, 2014), and (3) the computation of the difference between these two statistic values. We performed 1,000 such randomizations to obtain an estimate of the distribution of this difference under the null hypothesis. The analysis was performed from 0 ms to 350 ms after distractor onset, over 10 ms windows moving by step of 5 ms. To correct for multiple tests in the time dimension, a Bonferroni correction would have been over-conservative because it assumes that the data for each test are independent and does not take into account the temporal correlation of EEG data (Manly et al., 1986). To increase sensitivity while taking into account multiple testing in the time dimension, we used a randomization procedure proposed by Blair and Karniski (1993). For each permutation of the dataset, the maximum number of consecutive significant time-samples in the entire 0–350 ms time-window was computed. Over all the permutations, we thus obtained the distribution of this maximum number under the null-hypothesis. In this distribution, the 95th percentile corresponds to the maximum number of consecutive significant samples one can obtain by chance with a risk of 5%. We required all the effects to last at least this number of samples. This procedure was run separately for each channel. As the three electrodes were tested independently, a Bonferroni correction was applied on computed p-values to correct for family-wise errors (Weisstein, 2004).

Correlations across distracting sounds between ratings for arousal and valence and the median across participants of the mean amplitude of each of the ERPs mentioned below were conducted using Bayesian multiple linear regression. In our linear models, despite a correlation between arousal and valence ratings, the collinearity is not excessive with a Variance Inflation Factor (VIF) equal to 1.29, under the commonly accepted cutoff value of 5 or 10 (O’brien, 2007). Therefore, multiple linear regression with ratings for arousal and valence as regressors could be conducted. Tests were applied to each transient ERP at the electrode of their maximum: the N1 mean amplitude in the 80–120 ms window at Cz electrode, the P2 mean amplitude in the 140–180 ms window at Cz electrode, the early-P3 mean amplitude in the 200–260 ms window at Cz electrode, and the late-P3 mean amplitude in the 300–340 ms window at Fz and Pz electrodes. We report the Bayesian Factor (BF), the multiple R-squared for the results of the multiple linear regression and the partial R-squared for arousal and valence ratings, separately. Detailed output of the Bayesian regressions are reported in the Table 2.

**Table 2:** Results of Bayesian multiple linear regression on the amplitude of the N1, P2, early-P3 and late-P3 (frontal and parietal components) across the distracting sounds. The predictive variables were the arousal ratings and the valence ratings of the distracting sounds. The following outputs are specified: P(M)=prior model probability, P(M|data)=posterior model probability, BF_M_=change from prior to posterior model, BF_10_=Bayes Factor against the null model. The null model includes the subjects factor. For two-way Bayesian repeated-measure ANOVA, the following outputs are also specified: P(incl)=prior inclusion probability, P(incl|data)=posterior inclusion probability, BF_inclusion_=change from prior to posterior inclusion probability.

| N1 |  |  |  |  | Frontal late-P3 |  |  |  |  |
| --- | --- | --- | --- | --- | --- | --- | --- | --- | --- |
| Models | P(M) | P(M\|data) | BF<sub>M</sub> | BF<sub>10</sub> | Models | P(M) | P(M\|data) | BF<sub>M</sub> | BF<sub>10</sub> |
| Null model | 0.25 | 0.40 | 2.03 | 1.00 | Null model | 0.25 | 0.31 | 1.32 | 1.00 |
| Arousal | 0.25 | 0.16 | 0.55 | 0.39 | Arousal | 0.25 | 0.31 | 1.38 | 1.03 |
| Valence | 0.25 | 0.30 | 1.31 | 0.75 | Valence | 0.25 | 0.22 | 0.84 | 0.71 |
| Arousal + Valence | 0.25 | 0.14 | 0.47 | 0.34 | Arousal + Valence | 0.25 | 0.16 | 0.58 | 0.53 |
| Effects | P(incl) | P(incl\|data) | BF<sub>inclusion</sub> |  | Effects | P(incl) | P(incl\|data) | BF<sub>inclusion</sub> |  |
| Arousal | 0.50 | 0.29 | 0.41 |  | Arousal | 0.50 | 0.48 | 0.91 |  |
| Valence | 0.50 | 0.44 | 0.79 |  | Valence | 0.50 | 0.38 | 0.61 |  |

| P2 |  |  |  |  | Parietal late-P3 |  |  |  |  |
| --- | --- | --- | --- | --- | --- | --- | --- | --- | --- |
| Models | P(M) | P(M\|data) | BF<sub>M</sub> | BF<sub>10</sub> | Models | P(M) | P(M\|data) | BF<sub>M</sub> | BF<sub>10</sub> |
| Null model | 0.25 | 0.04 | 0.11 | 1.00 | Null model | 0.25 | 0.32 | 1.43 | 1.00 |
| Arousal | 0.25 | 0.51 | 3.14 | 0.34 | Arousal | 0.25 | 0.34 | 1.54 | 1.05 |
| Valence | 0.25 | 0.44 | 2.36 | 12.09 | Valence | 0.25 | 0.18 | 0.67 | 0.57 |
| Arousal + Valence | 0.25 | 0.01 | 0.04 | 14.06 | Arousal + Valence | 0.25 | 0.16 | 0.55 | 0.48 |
| Effects | P(incl) | P(incl\|data) | BF<sub>inclusion</sub> |  | Effects | P(incl) | P(incl\|data) | BF<sub>inclusion</sub> |  |
| Arousal | 0.50 | 0.52 | 1.10 |  | Arousal | 0.50 | 0.50 | 0.98 |  |
| Valence | 0.50 | 0.95 | 19.46 |  | Valence | 0.50 | 0.34 | 0.51 |  |

**Table 2:** Results of Bayesian multiple linear regression on the amplitude of the N1, P2, early-P3 and late-P3 (frontal and parietal components) across the distracting sounds.
| Early-P3 |  |  |  |  |
| --- | --- | --- | --- | --- |
| Models | P(M) | P(M\|data) | BF<sub>M</sub> | BF<sub>10</sub> |
| Null model | 0.25 | 4.95E-04 | 1.48E-03 | 1.00 |
| Arousal | 0.25 | 0.77 | 10.11 | 1558.71 |
| Valence | 0.25 | 3.35E-04 | 1.01E-3 | 0.06 |
| Arousal + Valence | 0.25 | 0.23 | 0.89 | 460.87 |
| Effects | P(incl) | P(incl\|data) | BF<sub>inclusion</sub> |  |
| Arousal | 0.50 | 0.99 | 1203.67 |  |
| Valence | 0.50 | 0.26 | 0.30 |  |

### 4.2 Results

#### 4.2.1 Properties of distracting sounds

The median arousal rating of all distracting sounds was 5.5. 15 high-arousing (HA) sounds (Mean ± SD = 6.0 ± 0.4) were found significantly more arousing than the 15 low-arousing (LA) sounds (Mean ± SD = 5.1 ± 0.3) (Mann-Whitney U test: p<0.001). We found a trend for a significant difference in valence ratings of HA and LA (HA: 4.7 ± 0.4, LA: 5.0 ± 0.5, Mann-Whitney U test: p=0.056). Ratings for arousal and valence were negatively (slope=-0.60) and significantly correlated (linear regression: F_1,28_=7.8, p=0.009, R^2^=0.21): negative sounds tended to be rated more arousing than neutral and positive sounds. Overall, sounds were rather neutral in valence (min=4.0, max=5.8, median=4.9).

No significant difference in mean pitch, mean roughness, mean brightness and attack leap was found between HA and LA sounds (Mann-Whitney U test: all p-values > 0.35).

#### 4.2.2 Percentage of correct responses (Table 1)

Participants correctly performed the detection task in 93.1% ± 0.9% of the trials. The remaining trials were either missed trials (1.5%) or trials with FAs (5.4% ± 0.9%).

The best model was the one with only the main effect of DISTRACTOR (BF_10_=5.0·10^16^). There was positive evidence against a specific effect of AROUSAL (BF_inclusion_=0.1) on RTs. There was no evidence in favor of the DISTRACTOR by AROUSAL interaction (BF_inclusion_=0.4), but there was decisive evidence of a specific effect of DISTRACTOR (BF_inclusion_=5.0·10^16^). Participants were less accurate in trials including distracting sounds (90.0 ± 1.4%) than in trials with no distracting sounds (95.6 ± 0.8%).

#### 4.2.3 Reaction times (Fig. 3, Table 1)

The best model was the one with only the main effect of DISTRACTOR (BF_10_=6.3·10^48^). There was positive evidence against a specific effect of AROUSAL on RTs (BF_inclusion_ =0.2) and a decisive specific effect of DISTRACTOR on RTs (BF_inclusion_ =6.4·10^48^).

Post-hoc analysis of the DISTRACTOR main effect showed that participants were faster in the DIS1 condition than in the NoDIS condition (BF_10_=5.9·10^23^) and than in the DIS3 condition (BF_10_=1.6·10^18^).

#### 4.2.4 EEG response to distracting sounds (Fig. 4)

**Fig. 4.**
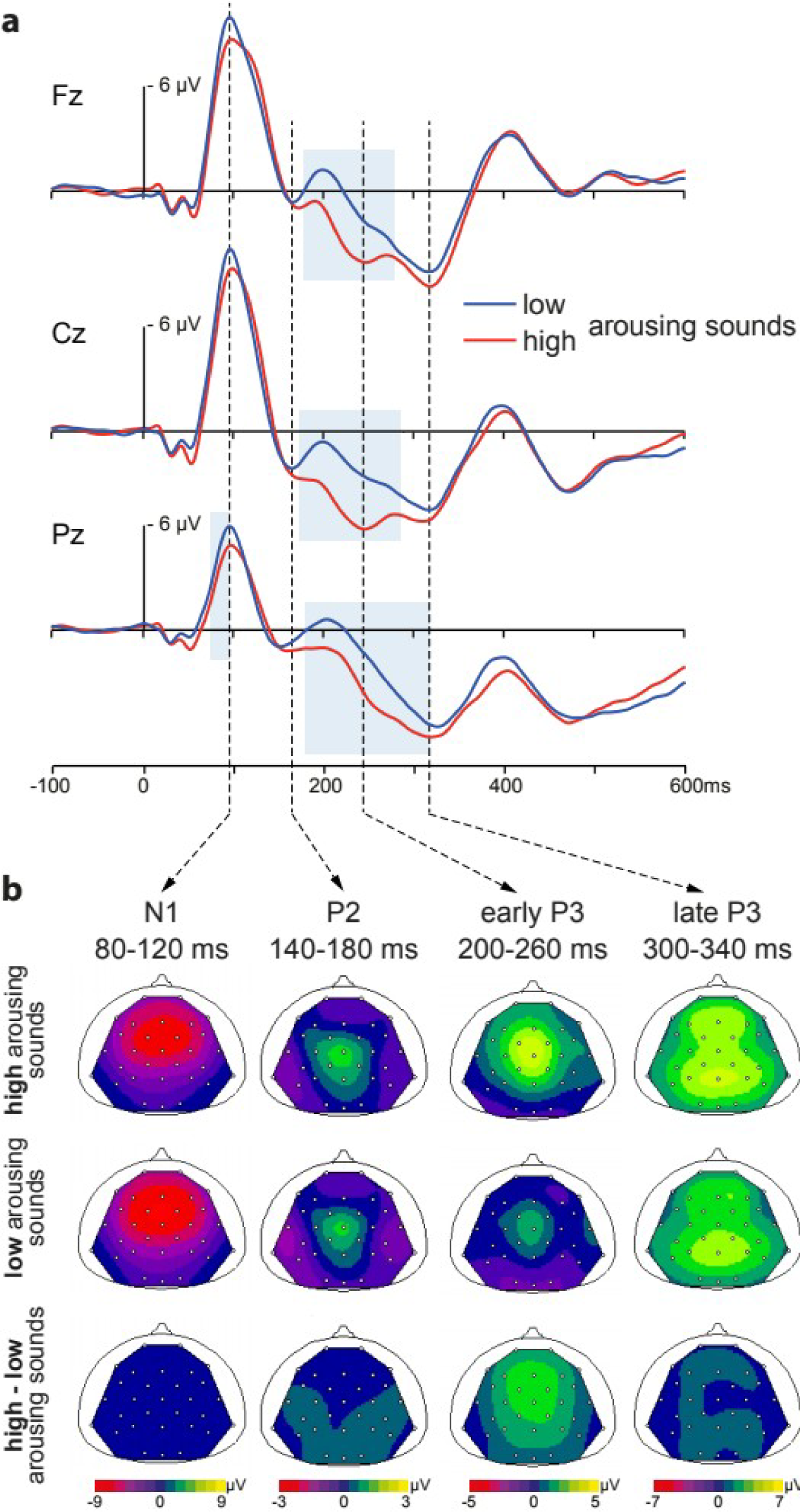
ERPs to distracting sounds. **a.** Mean ERPs (after subtraction of surrogate ERPs in the NoDIS condition) at the Fz, Cz and Pz electrodes as a function of the arousal category (high-arousing or low-arousing). The blue square corresponds to latencies for which a significant effect of the arousal category is observed (p < 0.05). **b.** From left to right, scalp topographies (top views) of the N1, P2, early-P3 and late-P3, in the high and low arousing categories and their difference, in the 80-120, 140-180, 200-260 and 300-340 ms time-windows after distractor onset, respectively.

In response to distracting sounds, the fronto-central N1 response (~100 ms) was followed by a small fronto-central P2 response (~160 ms), and a P3 complex that could be dissociated in two parts: an early-P3 (~240 ms) with a fronto-central distribution, and a late-P3 (~320 ms) with frontal and parietal components.

At the frontal electrode, the amplitude of the ERPs was significantly more positive for HA distractors from 180 ms to 280 ms (p-values ranging from <0.001 to 0.027), with the maximal difference at 221 ms. At the central electrode, the amplitude of the ERPs was significantly more positive for HA distractors from 175 ms to 290 ms (p-values ranging from <0.001 to 0.045), with the maximal difference at 223 ms. At the parietal electrode, the amplitude of the ERPs was significantly more positive for HA distractors from 80 ms to 100 ms (p-values ranging from <0.001 to 0.045) and from 180 ms to 300 ms (p-values ranging from <0.001 to 0.039), with the maximal difference at 253 ms. Please note that the amplitude of the ERPs at the parietal electrode were significant between 300 and 335 ms before Bonferroni correction but failed to reach significance after correction.

#### 4.2.5 Relationships between EEG responses to distracting sounds and sound ratings (Fig. 5, Table 2)

**Fig. 5.**
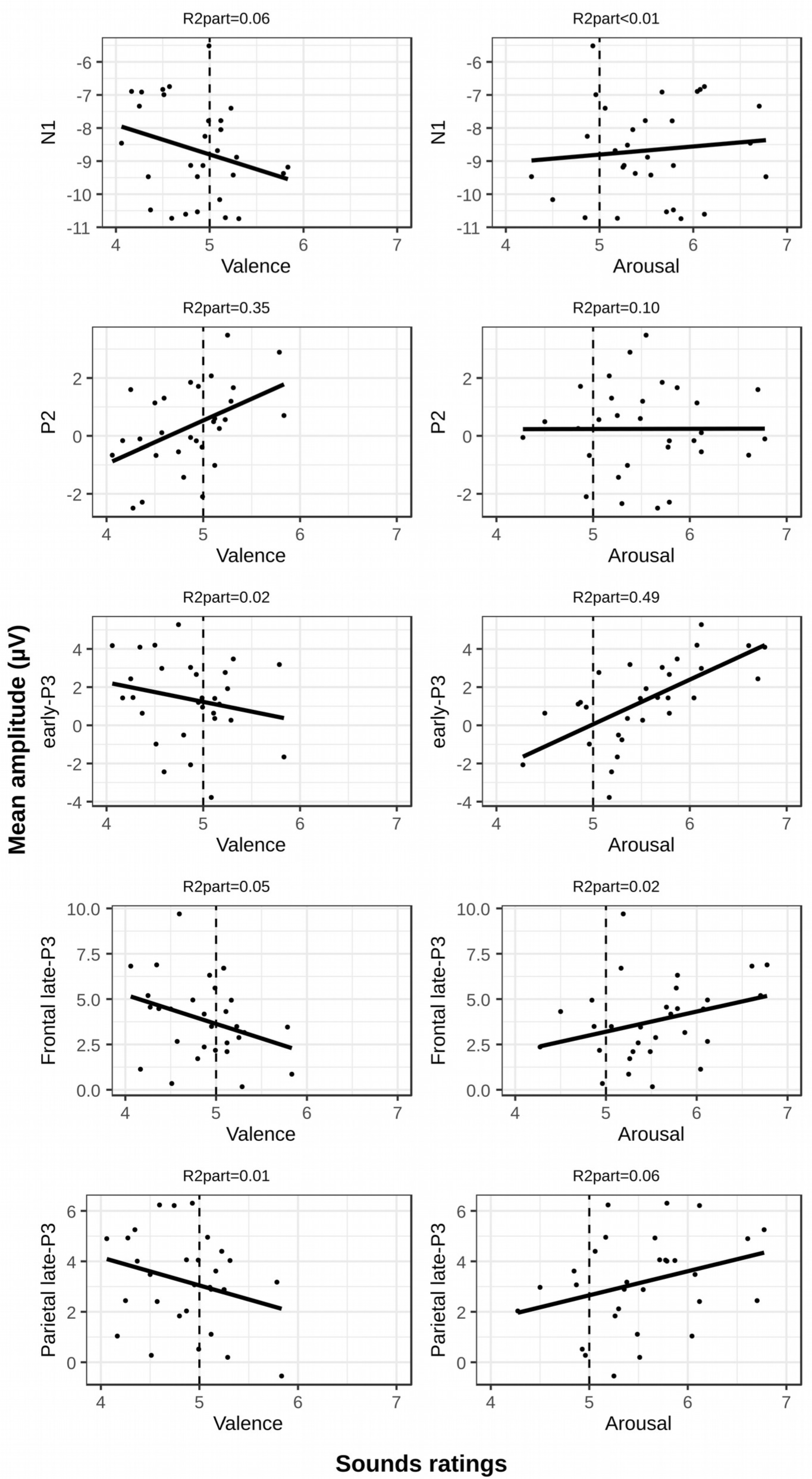
Scatterplots of the mean amplitude of the N1, P2, early-P3 and late-P3 across subjects for each distracting sound as a function of the arousing ratings (left) or the valence ratings (right). Amplitudes were measured at the Cz electrode group for the N1, P2 and early-P3 and at the Fz and Pz electrodes for the late-P3. A dashed vertical bar separates sounds rated as negative (below 5 out of 10) and sounds rated as positive (above 5 out of 10). We report partial R-squared (calculated as described in paragraph 4.1.7).

There was no evidence of an effect of arousal ratings (BF_10_=0.4), nor valence ratings (BF_10_=0.8) on the amplitude of the N1 (R^2^=0.05).

The best model of the P2 amplitude was the one including both effects of arousal and valence ratings (BF_10_=14.1) but it was only 1.2 times better than the model including only the effect of valence ratings (BF_10_=12.1). There was strong evidence of the specific effect of valence ratings (BF_inclusion_=19.5), but no evidence of a specific effect of arousal ratings (BF_inclusion_=1.1). The more positive the valence, the larger the P2 (R^2^_valence_=0.35).

The best model of the early-P3 amplitude was the one only including the effect of arousal ratings (BF_10_=1558.7). It was only 3.4 times better than the model including both arousal and valence rating effects (BF_10_=460.9). There was decisive evidence of the specific effect of arousal ratings (BF_inclusion_=1203.7), but there was positive evidence against a specific effect of valence ratings (BF_inclusion_=0.3). The more arousing, the larger the early-P3 (R^2^_arousal_=0.49).

There was no evidence of an effect of arousal ratings (BF_10_=1.0), nor of valence ratings (BF_10_=0.7), nor of both ratings (BF_10_=0.5) on the amplitude of the frontal late-P3 (R^2^=0.017).

There was no evidence of an effect of arousal ratings (BF_10_=1.1), nor of valence ratings (BF_10_=0.6), nor of both ratings (BF_10_=0.5) on the amplitude of the parietal late-P3 (R^2^=0.061).

### 4.3 Discussion

#### 4.3.1 Behavioral results

We found that the behavioral facilitation effect was not modulated according to the arousing content of the distraction sounds. The absence of an effect may be explained by the fact that in our study, high-arousing and low-arousing distractors were quite close in their mean arousal rating. This result contrasts with a recent audio-visual oddball study in which it has been reported that negative arousing novel sounds can trigger a smaller increase in reaction times compared to neutral novel sounds (Max et al., 2015). The authors interpreted this facilitation effect as “an unspecific benefit of emotional arousal”.

#### 4.3.2 Influence of the arousal and valence contents on ERPs

In the present study, in response to distracting sounds, two phases of the P3 response could be separated: an early phase peaking fronto-centrally between 200 and 260 ms (early-P3) and a late phase peaking frontally and parietally between 300 and 340 ms (late-P3) in agreement with previous studies (Escera et al., 1998; Escera and Corral, 2007; Yago et al., 2003).

We assessed the impact of the arousing value of distracting sounds on these electrophysiological responses: high-arousing distracting sounds elicited a larger early-P3. The difference between responses to high-arousing and low-arousing sounds was maximal at the latency of the early-P3 at fronto-central electrodes. The finding of increased early and late P3 to high arousing sounds is quite consistent with previous studies in the literature. In the auditory modality, both the early-P3 and the late-P3 have been found enhanced after high-arousing negative sounds compared to less-arousing neutral novel sounds (Widmann et al., 2018). In the visual modality, high-arousing novels trigger larger P3a waves than low-arousing novels regardless of their valence (Olofsson and Polich, 2007; Rozenkrants and Polich, 2008). However, another study showed that only the P3b (and not the P3a) was enhanced when comparing responses to negative/positive high-arousing novel pictures compared to low-arousing neutral pictures (Delplanque et al., 2005).

The distinction between arousal and emotional effects on the P3 responses has been often ignored since previous studies compared responses to high-arousing negative and/or positive stimuli, to less-arousing neutral ones, leading to debatable conclusions as valence and arousal can interact (Cuthbert et al., 2000; Delplanque et al., 2006; Domínguez-Borràs et al., 2008; Olofsson and Polich, 2007; Widmann et al., 2018). Here, we aimed at disentangling the effects of the arousing content of distracting sounds from the effects of their emotional content. First, we used distracting sounds that were rather emotionally neutral. Second, multiple regression analysis indicated strong to decisive evidence that the ERP correlated with arousal ratings, and not with the valence ratings, at the latencies of the early-P3: the more arousing the sound is, the larger the early-P3. Moreover, at the latencies of the late-P3, no evidence of a correlation with the arousal ratings, nor with the valence ratings, could be found. Finally, the valence ratings were only found to correlate with ERP amplitude at the latencies of the P2, which amplitude was enhanced for negative sounds. This finding is consistent with previous studies in the visual modality which report enhanced P2 amplitude to disgusting and fearful stimuli compared to neutral ones (Carretié et al., 2011, 2004). The P2 remains one of the least investigated ERP in the auditory modality: very little is known about its functional significance (Crowley and Colrain, 2004). In the present study, it has proven to be sensitive to the valence of distracting sounds but not to their arousing value.

Taken together, these results suggest that the amplitude of the early-P3, only, is strongly related to the arousal rating of the distracting sounds.

## 5. General discussion

### 5.1 Distracting sounds and arousal

We showed that “distracting” sounds can trigger a facilitation effect on behavior when the distractor-target delay is superior to 300 ms. This facilitation effect was found not affected by the sensory modality of the target and to remain stable for at least 2 seconds, suggesting that salient task-irrelevant sounds can increase general reactivity to any incoming stimulus.

All these results converge towards the idea that distracting sounds, on top of the behavioral costs related to attentional capture, can trigger a burst of arousal, which can result in a behavioral benefit. Phasic increases of arousal correspond to a state of enhanced physiological reactivity that leads to a condition of increased unspecific alertness and improved readiness to respond, which enables better performances. In tasks with low attentional demands, distracting sounds would temporarily increase arousal to a more optimal level enhancing reactivity during the task. This is consistent with a model of the orienting response which comprises an arousal component (Näätänen, 1992). Few studies investigating distraction have taken into account the arousal component provided by distracting sounds. In audio-visual oddball paradigms, distraction is measured as the difference in reaction time to targets preceded by a novel/deviant sounds compared to targets following a standard sound (Escera et al., 2000; Escera and Corral, 2007). It is important to note that since novel sounds can trigger both an attentional capture and an increase in phasic arousal, the behavioral costs and benefits resulting from each process, respectively, are both contributing to the so-called distraction measure in oddball paradigms.

In summary, distracting sounds can be followed by an increase in phasic arousal and, depending on the sound properties and the participant’s current level of tonic arousal, can lead to a substantial behavioral benefit. The effects of the arousal burst persist for a couple of seconds but decline with time.

### 5.2 The role of the fronto-central P3a in phasic arousal

The P3a has been frequently associated with distraction as its elicitation often coincided with impaired performances (Berti and Schröger, 2003; Escera et al., 2003; SanMiguel et al., 2008; Schröger and Wolff, 1998). P3a amplitude and the increase of RT due to deviant sounds is also moderately correlated (Berti et al., 2004). Furthermore, in an oddball paradigm, if deviant sounds become task-relevant, the P3a decreases and performances improve compared to task-irrelevant deviant sounds (Hölig and Berti, 2010). These results have been interpreted as an implication of the P3a in both involuntary and voluntary attention switching.

However, in the present study, the elicitation of both P3a phases was associated with improved performances (early distractors) but also with unchanged performances (late distractors). This is consistent with previous studies reporting a dissociation between the P3a and observable behavioral distraction (Rinne et al., 2006; SanMiguel et al., 2010b; Wetzel et al., 2013). As demonstrated earlier, salient task-irrelevant sounds trigger an orienting response which comprises both attentional capture and arousal components. These sounds can lead to improved, stable or impaired performances depending on task demands, distractor-target delays, the psychological state of the participant, etc. In other words, distracting sounds elicit both components of the orienting response, the final behavioral outcome depend on the balance between benefits and costs related to these two components. Therefore, linking the P3a solely to distraction may be reductive. In the light of our results and previous findings, the P3a can be seen as a more complex response which encompasses both components of the orienting response as it has been proposed by San Miguel and colleagues (2010).

In this line, previous results suggest that the early-P3 is not an adequate marker of attention reorientation. Contrary to the late-P3, the amplitude of the early-P3 is unchanged in a passive oddball paradigm compared to an active paradigm in which sounds precede the onset of visual targets (Escera et al., 2000, 1998). The authors concluded that as the early-P3 was insensitive to attentional manipulations, it was unlikely to reflect attentional reorientation. Sounds with personal significance (such as one’s own first name spoken by a familiar voice or one’s own ringtone) are expected to trigger stronger attentional capture but unlike the late-P3, the early-P3 is impervious to personal significance (Holeckova et al., 2006; Roye et al., 2007). In contrast to the late-P3, the early-P3 is observable only for high magnitude deviant sounds and not for low-magnitude deviant sounds even if both result in behavioral distraction (Horváth et al., 2011). In clinical studies, the early-P3 does not appear to be a reliable index of distraction. In a study using an auditory oddball in children with ADHD, enhanced distractibility of the children was associated with a smaller early-P3 to novel sounds but also with a larger late-P3 (Gumenyuk et al., 2005). Similar results were observable among schizophrenic patients who displayed a smaller early-P3 to novel sounds in association with an increased behavioral distraction (Cortiñas et al., 2008). Different interpretations of the functional role of the early-P3 have emerged: the early-P3 was proposed to reflect an alerting process governing the direction of the attentional move (Ceponiene et al., 2004), a stimulus-specific process (Horváth et al., 2011), or the violation of the regularity registered by the automatic deviance detection system (Escera et al., 2000, 1998).

In the present study, we found that the amplitude of the early-P3 is strongly related to the arousal rating of the distracting sounds. Therefore, we propose that the early-P3 is an index of the phasic arousal increase triggered by unexpected salient sounds. In agreement with previous articles, we hypothesize that the attentional capture is more likely to be reflected by the late-P3 as already proposed in several previous articles (Barry and Rushby, 2006; Bidet-Caulet et al., 2015; Escera et al., 2000, 1998; Roye et al., 2007).

Nomenclature of the P3-complex has been inconsistent throughout the literature. In active and passive oddball paradigms, infrequent target or non-target deviant sounds trigger a fronto-central P3a (Escera et al., 1998, 2000; Schröger and Wolff, 1998; Squires et al., 1975). In “novelty oddball” paradigms, it has been observed that novel environmental sounds were followed by an ERP coined as “novelty-P3” (Courchesne et al., 1975; Friedman et al., 2001; Gaeta et al., 2003). In an influential study, the authors described the novelty-P3 and the P3a as indistinguishable and that the two labels could be used interchangeably (Simons et al., 2001). This result has been widely accepted in the literature (Polich, 2007) but it has not prevented both nomenclatures to persist in subsequent articles (Escera and Corral, 2007; SanMiguel et al., 2010b). Moreover, both the P3a and the novelty-P3 have been found to comprise two phases: a fronto-central early phase peaking around 235 ms and a fronto-parietal late phase peaking around 320 ms (Escera et al., 1998; Escera and Corral, 2007; Yago et al., 2003). A recent study has run a Principal Component Analysis (PCA) compiling data from different oddball paradigms (Barry et al., 2016). Based on latency, topography and conditions of elicitation, the authors suggest that the early P3a reflects the genuine P3a and that the late-P3a reflects the genuine novelty-P3. In numerous articles, a monophasic fronto-central ERP peaking around 240 ms is referred to as “P3a”: this would be equivalent to the early-P3 observed here and in previous studies.

In the light of previous and present findings, it seems important to consider, in future studies, the different phases of the P3 complex in response to novel or distracting sounds. The investigation of their link with the distinct arousal and attentional components of the orienting response should provide new insight on the contradictory effect observed at the behavioral level.

### 5.3 Relationships between the fronto-central P3a and arousal

*From now on, “P3a” will refer specifically to the positive fronto-central ERP peaking around 240 ms, named early-P3 in the present study.*

Both P3a and arousal have been linked to the locus coeruleus-norepinephrine (LC-NE) system in the literature. The locus-coeruleus is a brainstem neuromodulatory system that sends noradrenergic innervation to all cortical regions and to the amygdala and notably influences the cortical control of attention through arousal (for a review, see Sara and Bouret, 2012). It has been proposed that the interaction between the NE system and the prefrontal cortex is crucial for attentional regulation (Arnsten, 1998). Excessive release of norepinephrine coincides with amplified responses to distracting stimuli and degraded task performance in monkeys (Rajkowski et al., 1997). The LC-NE would be closely linked to the orienting response: unexpected and/or arousing stimuli trigger a phasic activation of the LC-NE which consequently facilitates sensory processing (Aston-Jones and Cohen, 2005; Nieuwenhuis et al., 2005). P3a response and phasic activation of the LC-NE system share common antecedent conditions as both of them are elicited by motivationally significant stimuli such as novel stimuli in oddball tasks. In a passive auditory oddball study in monkeys, surgical lesions of the locus coeruleus were associated to decreased P3a-like response to deviant sounds (Pineda et al., 1989). It has been proposed that the LC-NE system phasic activation mediates both the P3a response and the behavioral facilitation effect following novel stimuli but such direct evidences in humans are still lacking (Schomaker and Meeter, 2015).

However, indirect evidences have accumulated. Autonomic components of the orienting responses such as the skin conductance response (SCR) and the pupil dilation response (PDR) are concomitant with the elicitation of the P3a and both would reflect the coactivation of the LC-NE system and the peripheral sympathetic nervous system (for a review, see Nieuwenhuis et al., 2011). In a recent oddball study, comodulation of the PDR response and the early-and late-P3a have been reported: both were enhanced in concert by high-arousing negative novel sounds compared to low-arousing neutral novel sounds (Widmann et al., 2018). However in another oddball study, at the single trial level, P3a did not correlate with the PDR amplitude (Kamp and Donchin, 2015). In oddball studies, a tendency towards a larger P3a has been observed in trials in which a SCR has been elicited (Lyytinen et al., 1992; Marinkovic et al., 2001) but other studies fail to link the SCR to any of the ERPs triggered by novel sounds (Barry et al., 2013; Rushby and Barry, 2009). In a recent auditory oddball study, large SCR to novel sounds were associated to large late-P3 amplitudes but no such association could be drawn with the early-P3 (Berti et al., 2017). Studies using drugs to modulate noradrenergic activity can also be useful to understand the links between the P3a and the LC-NE system. A drug facilitating noradrenergic neurotransmission has been found to increase specifically the P3a in an auditory oddball, and had no effect on the P3b (Missonnier et al., 1999). Clonidine, a drug which attenuates baseline noradrenergic activity, has been found to decrease the amplitude of the P3a (but to increase P3b) during an auditory oddball task (Brown et al., 2015). Further research will be necessary to corroborate the hypothesis of a causal link between the phasic increase of physiological arousal and the fronto-central early-P3 response after unexpected salient stimuli.

### 5.4 An arousal model

This model proposes that the fronto-central P3a reflect the phasic increase in arousal triggered by novel or distracting sounds. This alternative interpretation of the processes underlying the fronto-central P3a is in line with the Näätänen model of the orienting response including an arousal and an attentional components, and with a link between the P3a and the LC-NE system (see previous paragraph). In monkeys, the locus coeruleus is activated around 100 ms after stimulus onset (Aston-Jones et al., 1994; Rajkowski et al., 1994). It is imaginable that the P3a elicitation (whose peak is around 240 ms after stimulus onset) is the result of the general increase of cortical excitability following the LC-NE activation by novel stimuli.

This model is consistent with previous results and can help to shed light on them. First, it has been observed repeatedly that novel sounds trigger a larger fronto-central P3a than deviant sounds (e.g. Escera et al., 1998; Fabiani and Friedman, 1995; Gaeta et al., 2003; Spencer et al., 1999). This effect cannot be explained simply by deviance (both deviant and novel sounds deviate from the context of the standard sound sequence), nor by novelty (here understood as the association of rareness and unfamiliarity) because white noise distractors trigger a larger P3a than variable environmental novel sounds (Frank et al., 2012) and identical environmental sounds trigger a similar P3a than variable environmental novel sounds (Barkaszi et al., 2013). The difference in fronto-central P3a amplitude between novel and deviant sounds could, however, be explained by the fact that novel sounds are more salient due to their complexity and therefore more arousing, resulting in a steeper increase in phasic arousal and a larger fronto-central P3a. Second, one of the main characteristics of the P3a is to quickly decrement with repeated exposure (Barry et al., 2016; Courchesne et al., 1975; Debener et al., 2002). This is compatible with the observations that the locus coeruleus neural response to novel sounds or to novel environments quickly decreases after few presentations in rats (Hervé-Minvielle and Sara, 1995; Vankov et al., 1995). Therefore, the arousal model of the P3a fits the classic theory of the orienting response and can account for the main properties of the P3a that have been highlighted in the literature.

P3a has been widely considered as an index of distractibility and used as such in numerous clinical studies including for example healthy old adults (Getzmann et al., 2013), participants under the influence of alcohol (Marinkovic et al., 2001), patients with closed head injury (Kaipio et al., 2000), Alzheimer (Correa-Jaraba et al., 2018) or dyslexia (Rüsseler et al., 2002), healthy children (e.g. Gumenyuk et al., 2001; Wetzel and Schröger, 2007) and children with ASD (Ferri et al., 2003), depression (Lepistö et al., 2004) or ADHD (e.g. Gumenyuk et al., 2005; Keage et al., 2006; van Mourik et al., 2007). Misunderstanding the complexity of the P3a signal could lead to erroneous interpretations of data in clinical studies. An increased arousability (in the sense of a propensity to increased phasic arousal responses) could be mistaken for an enhanced distractibility. This arousal model of the P3a may help to make sense to seemingly paradoxical results such as the association of a reduced P3a with increased behavioral distraction (van Mourik et al., 2007; Keage et al., 2006) or opposite modulations of the early-and late-P3 (Cortiñas et al., 2008; Gumenyuk et al., 2005). It will be essential for further studies about distractibility to dissociate the two phases of the P3 complex to improve medical diagnosis.

## 6. Conclusions

In summary, the present findings show that unexpected salient sounds during a task trigger two distinct processes: a short-lived attentional capture and a more lingering phasic increase in arousal. Consequently, performances are conditioned to both facilitation by phasic arousal and impairment by attentional capture. Whether unexpected salient sounds will have detrimental or beneficial effects on performances partly depends on the design of the task (the sound-target delay in the present study but also the informational value or task-relevance of the sound, the task demand, etc.).

Unexpected salient sounds trigger a biphasic P3, we found that the amplitude of the early-P3 is strongly related to the arousal rating of the distracting sounds. Therefore, we propose that both the attention and arousal components of the orienting response are reflected in the P3 but that the phasic arousal response is more associated to the fronto-central early-P3, often named P3a in the literature.

## 8. Acknowledgments

We thank A. Garnier, L. Bottemanne, B. Chatard, B. Chen and R. Dhillon for their help in recruiting, testing subjects and for preliminary analyses. We thank P. Ruby for proofreading our manuscript and for her insights on the aforementioned results and A. Moulin and A. Caclin for their insights on statistical analyses.

## References

Andrés, P., Parmentier, F.B.R., Escera, C., 2006. The effect of age on involuntary capture of attention by irrelevant sounds: a test of the frontal hypothesis of aging. Neuropsychologia 44, 2564–2568. https://doi.org/10.1016/j.neuropsychologia.2006.05.005Arnsten, A.F.T., 1998. Catecholamine modulation of prefrontal cortical cognitive function. Trends in Cognitive Sciences 2, 436–447. https://doi.org/10.1016/S1364-6613(98)01240-6

Aston-Jones, G., Cohen, J.D., 2005. An integrative theory of locus coeruleus-norepinephrine function: adaptive gain and optimal performance. Annu. Rev. Neurosci. 28, 403–450. https://doi.org/10.1146/annurev.neuro.28.061604.135709

Aston-Jones, G., Rajkowski, J., Kubiak, P., Alexinsky, T., 1994. Locus coeruleus neurons in monkey are selectively activated by attended cues in a vigilance task. J. Neurosci. 14, 4467–4480. https://doi.org/10.1523/JNEUROSCI.14-07-04467.1994

Barkaszi, I., Czigler, I., Balázs, L., 2013. Stimulus complexity effects on the event-related potentials to task-irrelevant stimuli. Biological Psychology 94, 82–89. https://doi.org/10.1016/j.biopsycho.2013.05.007

Barry, R.J., MacDonald, B., De Blasio, F.M., Steiner, G.Z., 2013. Linking components of event-related potentials and autonomic measures of the orienting reflex. International Journal of Psychophysiology, Psychophysiology in Australasia - ASP conference - November 28-30 2012 89, 366–373. https://doi.org/10.1016/j.ijpsycho.2013.01.016

Barry, R.J., Rushby, J.A., 2006. An orienting reflex perspective on anteriorisation of the P3 of the event-related potential. Exp Brain Res 173, 539–545. https://doi.org/10.1007/s00221-006-0590-8

Barry, R.J., Steiner, G.Z., De Blasio, F.M., 2016. Reinstating the Novelty P3. Sci Rep 6. https://doi.org/10.1038/srep31200

Berti, S., 2013. The role of auditory transient and deviance processing in distraction of task performance: a combined behavioral and event-related brain potential study. Front Hum Neurosci 7. https://doi.org/10.3389/fnhum.2013.00352

Berti, S., Roeber, U., Schröger, E., 2004. Bottom-Up Influences on Working Memory: Behavioral and Electrophysiological Distraction Varies with Distractor Strength. Experimental Psychology 51, 249–257. https://doi.org/10.1027/1618-3169.51.4.249

Berti, S., Schröger, E., 2003. Working memory controls involuntary attention switching: evidence from an auditory distraction paradigm. European Journal of Neuroscience 17, 1119–1122. https://doi.org/10.1046/j.1460-9568.2003.02527.x

Berti, S., Vossel, G., Gamer, M., 2017. The Orienting Response in Healthy Aging: Novelty P3 Indicates No General Decline but Reduced Efficacy for Fast Stimulation Rates. Front Psychol 8. https://doi.org/10.3389/fpsyg.2017.01780

Bidet-Caulet, A., Bottemanne, L., Fonteneau, C., Giard, M.-H., Bertrand, O., 2015. Brain dynamics of distractibility: interaction between top-down and bottom-up mechanisms of auditory attention. Brain Topogr 28, 423–436. https://doi.org/10.1007/s10548-014-0354-x

Blair, R.C., Karniski, W., 1993. An alternative method for significance testing of waveform difference potentials. Psychophysiology 30, 518–524. https://doi.org/10.1111/j.1469-8986.1993.tb02075.x

Bradley, M.M., Lang, P.J., 1994. Measuring emotion: the Self-Assessment Manikin and the Semantic Differential. J Behav Ther Exp Psychiatry 25, 49–59.

Brown, S.B.R.E., van der Wee, N.J.A., van Noorden, M.S., Giltay, E.J., Nieuwenhuis, S., 2015. Noradrenergic and cholinergic modulation of late ERP responses to deviant stimuli. Psychophysiology 52, 1620–1631. https://doi.org/10.1111/psyp.12544

Carretié, L., Hinojosa, J.A., Martín-Loeches, M., Mercado, F., Tapia, M., 2004. Automatic attention to emotional stimuli: neural correlates. Hum Brain Mapp 22, 290–299. https://doi.org/10.1002/hbm.20037

Carretié, L., Ruiz-Padial, E., López-Martín, S., Albert, J., 2011. Decomposing unpleasantness: differential exogenous attention to disgusting and fearful stimuli. Biol Psychol 86, 247–253. https://doi.org/10.1016/j.biopsycho.2010.12.005

Ceponiene, R., Lepistö, T., Soininen, M., Aronen, E., Alku, P., Näätänen, R., 2004. Event-related potentials associated with sound discrimination versus novelty detection in children. Psychophysiology 41, 130–141. https://doi.org/10.1111/j.1469-8986.2003.00138.x

Correa-Jaraba, K.S., Lindín, M., Díaz, F., 2018. Increased Amplitude of the P3a ERP Component as a Neurocognitive Marker for Differentiating Amnestic Subtypes of Mild Cognitive Impairment. Front Aging Neurosci 10. https://doi.org/10.3389/fnagi.2018.00019

Cortiñas, M., Corral, M.-J., Garrido, G., Garolera, M., Pajares, M., Escera, C., 2008. Reduced novelty-P3 associated with increased behavioral distractibility in schizophrenia. Biological Psychology 78, 253–260. https://doi.org/10.1016/j.biopsycho.2008.03.011

Coull, J.T., 1998. Neural correlates of attention and arousal: insights from electrophysiology, functional neuroimaging and psychopharmacology. Progress in Neurobiology 55, 343–361. https://doi.org/10.1016/S0301-0082(98)00011-2

Courchesne, E., Hillyard, S.A., Galambos, R., 1975. Stimulus novelty, task relevance and the visual evoked potential in man. Electroencephalography and Clinical Neurophysiology 39, 131–143. https://doi.org/10.1016/0013-4694(75)90003-6

Crowley, K.E., Colrain, I.M., 2004. A review of the evidence for P2 being an independent component process: age, sleep and modality. Clinical Neurophysiology 115, 732–744. https://doi.org/10.1016/j.clinph.2003.11.021

Cuthbert, B.N., Schupp, H.T., Bradley, M.M., Birbaumer, N., Lang, P.J., 2000. Brain potentials in affective picture processing: covariation with autonomic arousal and affective report. Biological Psychology 52, 95–111. https://doi.org/10.1016/S0301-0511(99)00044-7

Debener, S., Kranczioch, C., Herrmann, C.S., Engel, A.K., 2002. Auditory novelty oddball allows reliable distinction of top-down and bottom–up processes of attention. International Journal of Psychophysiology 46, 77–84. https://doi.org/10.1016/S0167-8760(02)00072-7

Delplanque, S., Silvert, L., Hot, P., Rigoulot, S., Sequeira, H., 2006. Arousal and valence effects on event-related P3a and P3b during emotional categorization. International Journal of Psychophysiology 60, 315–322. https://doi.org/10.1016/j.ijpsycho.2005.06.006

Delplanque, S., Silvert, L., Hot, P., Sequeira, H., 2005. Event-related P3a and P3b in response to unpredictable emotional stimuli. Biological Psychology 68, 107–120. https://doi.org/10.1016/j.biopsycho.2004.04.006

Domínguez-Borrás, J., Garcia-Garcia, M., Escera, C., 2008. Emotional context enhances auditory novelty processing: behavioural and electrophysiological evidence. The European Journal Of Neuroscience 28, 1199–1206. https://doi.org/10.1111/j.1460-9568.2008.06411.x

Edgington, E.S., 2014. Randomization Tests, in: Wiley StatsRef: Statistics Reference Online. American Cancer Society. https://doi.org/10.1002/9781118445112.stat01772

Escera, C., Alho, K., Schröger, E., Winkler, I., 2000. Involuntary attention and distractibility as evaluated with event-related brain potentials. Audiol. Neurootol. 5, 151–166. https://doi.org/13877

Escera, C., Alho, K., Winkler, I., Näätänen, R., 1998. Neural mechanisms of involuntary attention to acoustic novelty and change. Journal of Cognitive Neuroscience 590–604.

Escera, C., Corral, M. j., 2007. Role of Mismatch Negativity and Novelty-P3 in Involuntary Auditory Attention. Journal of Psychophysiology 21, 251–264. https://doi.org/10.1027/0269-8803.21.34.251

Escera, C., Yago, E., Corral, M.-J., Corbera, S., Nuñez, M.I., 2003. Attention capture by auditory significant stimuli: semantic analysis follows attention switching. Eur. J. Neurosci. 18, 2408–2412.

Ferri, R., Elia, M., Agarwal, N., Lanuzza, B., Musumeci, S.A., Pennisi, G., 2003. The mismatch negativity and the P3a components of the auditory event-related potentials in autistic low-functioning subjects. Clinical Neurophysiology 114, 1671–1680. https://doi.org/10.1016/S1388-2457(03)00153-6

Frank, D.W., Yee, R.B., Polich, J., 2012. P3a from white noise. International Journal of Psychophysiology 85, 236–241. https://doi.org/10.1016/j.ijpsycho.2012.04.005

Friedman, D., Cycowicz, Y.M., Gaeta, H., 2001. The novelty P3: an event-related brain potential (ERP) sign of the brain’s evaluation of novelty. Neuroscience & Biobehavioral Reviews 25, 355–373. https://doi.org/10.1016/S0149-7634(01)00019-7

Gaeta, H., Friedman, D., Hunt, G., 2003. Stimulus characteristics and task category dissociate the anterior and posterior aspects of the novelty P3. Psychophysiology 40, 198–208.

Getzmann, S., Gajewski, P.D., Falkenstein, M., 2013. Does age increase auditory distraction? Electrophysiological correlates of high and low performance in seniors. Neurobiology of Aging 34, 1952–1962. https://doi.org/10.1016/j.neurobiolaging.2013.02.014

Guillaume, A., Drake, C., Blancard, C., Chastres, V., Pellieux, L., 2004. How long does it take to identify everyday sounds.

Gumenyuk, V., Korzyukov, O., Escera, C., Hämäläinen, M., Huotilainen, M., Häyrinen, T., Oksanen, H., Näätänen, R., von Wendt, L., Alho, K., 2005. Electrophysiological evidence of enhanced distractibility in ADHD children. Neuroscience Letters 374, 212–217. https://doi.org/10.1016/j.neulet.2004.10.081

Hackley, S.A., 2009. The speeding of voluntary reaction by a warning signal. Psychophysiology 46, 225–233. https://doi.org/10.1111/j.1469-8986.2008.00716.x

Hackley, S.A., Valle-Inclán, F., 1999. Accessory stimulus effects on response selection: does arousal speed decision making? J Cogn Neurosci 11, 321–329.

Hackley, S.A., Valle-Inclán, F., 1998. Automatic alerting does not speed late motoric processes in a reaction-time task. Nature 391, 786–788. https://doi.org/10.1038/35849

Hervé-Minvielle, A., Sara, S.J., 1995. Rapid habituation of auditory responses of locus coeruleus cells in anaesthetized and awake rats. Neuroreport 6, 1363–1368.

Holeckova, I., Fischer, C., Giard, M.-H., Delpuech, C., Morlet, D., 2006. Brain responses to a subject’s own name uttered by a familiar voice. Brain Research 1082, 142–152. https://doi.org/10.1016/j.brainres.2006.01.089

Holig, C., Berti, S., 2010. To switch or not to switch: Brain potential indices of attentional control after task-relevant and task-irrelevant changes of stimulus features. Brain Research 1345, 164–175. https://doi.org/10.1016/j.brainres.2010.05.047

Horváth, J., Sussman, E., Winkler, I., Schröger, E., 2011. Preventing distraction: Assessing stimulus-specific and general effects of the predictive cueing of deviant auditory events. Biological Psychology 87, 35–48. https://doi.org/10.1016/j.biopsycho.2011.01.011

Kaipio, M.L., Cheour, M., Ceponiene, R., Ohman, J., Alku, P., Näätänen, R., 2000. Increased distractibility in closed head injury as revealed by event-related potentials. Neuroreport 11, 1463–1468.

Kamp, S.-M., Donchin, E., 2015. ERP and pupil responses to deviance in an oddball paradigm. Psychophysiol 52, 460–471. https://doi.org/10.1111/psyp.12378

Keage, H.A.D., Clark, C.R., Hermens, D.F., Kohn, M.R., Clarke, S., Williams, L.M., Crewther, D., Lamb, C., Gordon, E., 2006. Distractibility in AD/HD predominantly inattentive and combined subtypes: the P3a ERP component, heart rate and performance. J. Integr. Neurosci. 5, 139–158.

Lartillot, O., Toiviainen, P., Eerola, T., 2008. A Matlab Toolbox for Music Information Retrieval, in: Data Analysis, Machine Learning and Applications, Studies in Classification, Data Analysis, and Knowledge Organization. Springer, Berlin, Heidelberg, pp. 261–268. https://doi.org/10.1007/978-3-540-78246-9_31

Lee, M.D., Wagenmakers, E.-J., 2014. Bayesian Cognitive Modeling: A Practical Course. Cambridge University Press.

Lepistö, T., Soininen, M., Čeponien, R., Almqvist, F., Näätänen, R., Aronen, E.T., 2004. Auditory event-related potential indices of increased distractibility in children with major depression. Clinical Neurophysiology 115, 620–627. https://doi.org/10.1016/j.clinph.2003.10.020

Li, B., Parmentier, F.B.R., Zhang, M., 2013. Behavioral distraction by auditory deviance is mediated by the sound’s informational value. Evidence from an auditory discrimination task. Exp Psychol 60, 260–268. https://doi.org/10.1027/1618-3169/a000196

Ljungberg, J.K., R, B., Leiva, A., Vega, N., 2012. The informational constraints of behavioral distraction by unexpected sounds: The role of event information. Journal of Experimental Psychology: Learning, Memory, and Cognition 38, 1461–1468. https://doi.org/10.1037/a0028149

Lyytinen, H., Blomberg, A.-P., Näätänen, R., 1992. Event-Related Potentials and Autonomic Responses to a Change in Unattended Auditory Stimuli. Psychophysiology 29, 523–534. https://doi.org/10.1111/j.1469-8986.1992.tb02025.x

Manly, B.F., McAlevey, L., Stevens, D., 1986. A randomization procedure for comparing group means on multiple measurements. British Journal of Mathematical and Statistical Psychology 39, 183–189. https://doi.org/10.1111/j.2044-8317.1986.tb00855.x

Marcell, M.M., Borella, D., Greene, M., Kerr, E., Rogers, S., 2000. Confrontation Naming of Environmental Sounds. Journal of Clinical and Experimental Neuropsychology 22, 830–864. https://doi.org/10.1076/jcen.22.6.830.949

Marinkovic, K., Halgren, E., Maltzman, I., 2001. Arousal-related P3a to novel auditory stimuli is abolished by a moderately low alcohol dose. Alcohol Alcohol. 36, 529–539.

Max, C., Widmann, A., Kotz, S.A., Schröger, E., Wetzel, N., 2015. Distraction by Emotional Sounds: Disentangling Arousal Benefits and Orienting Costs. Emotion No Pagination Specified. https://doi.org/10.1037/a0039041

Missonnier, P., Ragot, R., Derouesné, C., Guez, D., Renault, B., 1999. Automatic attentional shifts induced by a noradrenergic drug in Alzheimer’s disease: evidence from evoked potentials. International Journal of Psychophysiology 33, 243–251. https://doi.org/10.1016/S0167-8760(99)00059-8

Näätänen, R., 1992. Attention and Brain Function. Psychology Press.

Näätänen, R., 1990. The role of attention in auditory information processing as revealed by event-related potentials and other brain measures of cognitive function. Behavioral and Brain Sciences 13, 201–233. https://doi.org/10.1017/S0140525X00078407

Näätänen, R., Picton, T., 1987. The N1 Wave of the Human Electric and Magnetic Response to Sound: A Review and an Analysis of the Component Structure. Psychophysiology 24, 375–425. https://doi.org/10.1111/j.1469-8986.1987.tb00311.x

Näätänen, R., Winkler, I., 1999. The concept of auditory stimulus representation in cognitive neuroscience. Psychological Bulletin 125, 826–859. https://doi.org/10.1037/0033-2909.125.6.826

Nieuwenhuis, S., Aston-Jones, G., Cohen, J.D., 2005. Decision making, the P3, and the locus coeruleus‐‐norepinephrine system. Psychological Bulletin 131, 510–532. https://doi.org/10.1037/0033-2909.131.4.510

Nieuwenhuis, S., de Geus, E.J., Aston-Jones, G., 2011. The anatomical and functional relationship between the P3 and autonomic components of the orienting response. Psychophysiology 48, 162–175. https://doi.org/10.1111/j.1469-8986.2010.01057.x

O’brien, R.M., 2007. A Caution Regarding Rules of Thumb for Variance Inflation Factors. Qual Quant 41, 673–690. https://doi.org/10.1007/s11135-006-9018-6

Olofsson, J.K., Polich, J., 2007. Affective visual event-related potentials: Arousal, repetition, and time-on-task. Biological Psychology 75, 101–108. https://doi.org/10.1016/j.biopsycho.2006.12.006

Parmentier, F.B.R., 2014. The cognitive determinants of behavioral distraction by deviant auditory stimuli: a review. Psychol Res 78, 321–338. https://doi.org/10.1007/s00426-013-0534-4

Parmentier, F.B.R., Elford, G., Escera, C., Andrés, P., Miguel, I.S., 2008. The cognitive locus of distraction by acoustic novelty in the cross-modal oddball task. Cognition 106, 408–432. https://doi.org/10.1016/j.cognition.2007.03.008

Parmentier, F.B.R., Elsley, J.V., Andrés, P., Barceló, F., 2011. Why are auditory novels distracting? Contrasting the roles of novelty, violation of expectation and stimulus change. Cognition 119, 374–380. https://doi.org/10.1016/j.cognition.2011.02.001

Parmentier, F.B.R., Elsley, J.V., Ljungberg, J.K., 2010. Behavioral distraction by auditory novelty is not only about novelty: The role of the distracter’s informational value. Cognition 115, 504–511. https://doi.org/10.1016/j.cognition.2010.03.002

Perrin, F., Pernier, J., Bertrand, O., Echallier, J.F., 1989. Spherical splines for scalp potential and current density mapping. Electroencephalography and Clinical Neurophysiology 72, 184–187. https://doi.org/10.1016/0013-4694(89)90180-6

Pineda, J.A., Foote, S.L., Neville, H.J., 1989. Effects of locus coeruleus lesions on auditory, long-latency, event-related potentials in monkey. J. Neurosci. 9, 81–93. https://doi.org/10.1523/JNEUROSCI.09-01-00081.1989

Polich, J., 2007. Updating P300: An integrative theory of P3a and P3b. Clinical Neurophysiology 118, 2128–2148. https://doi.org/10.1016/j.clinph.2007.04.019

Polich, J., Criado, J.R., 2006. Neuropsychology and neuropharmacology of P3a and P3b. International Journal of Psychophysiology, Models and Theories of Brain Function with Special Emphasis on Cognitive Processing 60, 172–185. https://doi.org/10.1016/j.ijpsycho.2005.12.012

Posner, M.I., Nissen, M.J., Klein, R.M., 1976. Visual dominance: an information-processing account of its origins and significance. Psychol Rev 83, 157–171.

Rajkowski, J., Kubiak, P., Aston-Jones, G., 1994. Locus coeruleus activity in monkey: Phasic and tonic changes are associated with altered vigilance. Brain Research Bulletin 35, 607–616. https://doi.org/10.1016/0361-9230(94)90175-9

Rajkowski, J., Kubiak, P., Ivanova, S., Aston-Jones, G., 1997. State-Related Activity, Reactivity of Locus Ceruleus Neurons in Behaving Monkeys, in: Goldstein, D.S., Eisenhofer, G., McCarty, R. (Eds.), Advances in Pharmacology. Academic Press, pp. 740–744. https://doi.org/10.1016/S1054-3589(08)60854-6

Ranganath, C., Rainer, G., 2003. Neural mechanisms for detecting and remembering novel events. Nat. Rev. Neurosci. 4, 193–202. https://doi.org/10.1038/nrn1052

Rinne, T., Särkkä, A., Degerman, A., Schröger, E., Alho, K., 2006. Two separate mechanisms underlie auditory change detection and involuntary control of attention. Brain Research 1077, 135–143. https://doi.org/10.1016/j.brainres.2006.01.043

Roye, A., Jacobsen, T., Schröger, E., 2007. Personal significance is encoded automatically by the human brain: an event-related potential study with ringtones. European Journal of Neuroscience 26, 784–790. https://doi.org/10.1111/j.1460-9568.2007.05685.x

Rozenkrants, B., Polich, J., 2008. Affective ERP processing in a visual oddball task: Arousal, valence, and gender. Clinical Neurophysiology 119, 2260–2265. https://doi.org/10.1016/j.clinph.2008.07.213

Rushby, J.A., Barry, R.J., 2009. Single-trial event-related potentials to significant stimuli. International Journal of Psychophysiology 74, 120–131. https://doi.org/10.1016/j.ijpsycho.2009.08.003

Rüsseler, J., Kowalczuk, J., Johannes, S., Wieringa, B.M., Münte, T.F., 2002. Cognitive brain potentials to novel acoustic stimuli in adult dyslexic readers. Dyslexia 8, 125–142. https://doi.org/10.1002/dys.221

SanMiguel, I., Corral, M.-J., Escera, C., 2008. When Loading Working Memory Reduces Distraction: Behavioral and Electrophysiological Evidence from an Auditory-Visual Distraction Paradigm. Journal of Cognitive Neuroscience 20, 1131–1145. https://doi.org/10.1162/jocn.2008.20078

SanMiguel, I., Linden, D., Escera, C., 2010a. Attention capture by novel sounds: Distraction versus facilitation. European Journal of Cognitive Psychology 22, 481–515. https://doi.org/10.1080/09541440902930994

SanMiguel, I., Morgan, H.M., Klein, C., Linden, D., Escera, C., 2010b. On the functional significance of Novelty-P3: Facilitation by unexpected novel sounds. Biological Psychology 83, 143–152. https://doi.org/10.1016/j.biopsycho.2009.11.012

Schomaker, J., Meeter, M., 2015. Short‐ and long-lasting consequences of novelty, deviance and surprise on brain and cognition. Neuroscience & Biobehavioral Reviews 55, 268–279. https://doi.org/10.1016/j.neubiorev.2015.05.002

Schomaker, J., Meeter, M., 2014. Facilitation of responses by task-irrelevant complex deviant stimuli. Acta Psychologica 148, 74–80. https://doi.org/10.1016/j.actpsy.2014.01.006

Schröger, E., 1996. A Neural Mechanism for Involuntary Attention Shifts to Changes in Auditory Stimulation. Journal of Cognitive Neuroscience 8, 527–539. https://doi.org/10.1162/jocn.1996.8.6.527

Schröger, E., Wolff, C., 1998. Behavioral and electrophysiological effects of task-irrelevant sound change: a new distraction paradigm. Cognitive Brain Research 7, 71–87. https://doi.org/10.1016/S0926-6410(98)00013-5

Simons, R.F., Graham, F.K., Miles, M.A., Chen, X., 2001. On the relationship of P3a and the Novelty-P3. Biological Psychology 56, 207–218. https://doi.org/10.1016/S0301-0511(01)00078-3

Squires, N.K., Squires, K.C., Hillyard, S.A., 1975. Two varieties of long-latency positive waves evoked by unpredictable auditory stimuli in man. Electroencephalogr Clin Neurophysiol 38, 387–401.

Sturm, W., Willmes, K., 2001. On the Functional Neuroanatomy of Intrinsic and Phasic Alertness. NeuroImage 14, S76–S84. https://doi.org/10.1006/nimg.2001.0839

Tona, K.-D., Murphy, P.R., Brown, S.B.R.E., Nieuwenhuis, S., 2016. The accessory stimulus effect is mediated by phasic arousal: A pupillometry study. Psychophysiology 53, 1108–1113. https://doi.org/10.1111/psyp.12653

Valls-Solé, J., Solé, A., Valldeoriola, F., Muñoz, E., Gonzalez, L.E., Tolosa, E.S., 1995. Reaction time and acoustic startle in normal human subjects. Neuroscience Letters 195, 97–100. https://doi.org/10.1016/0304-3940(94)11790-P

Vankov, A., Hervé-Minvielle, A., Sara, S.J., 1995. Response to Novelty and its Rapid Habituation in Locus Coeruleus Neurons of the Freely Exploring Rat. European Journal of Neuroscience 7, 1180–1187. https://doi.org/10.1111/j.1460-9568.1995.tb01108.x

Wagenmakers, E.-J., Love, J., Marsman, M., Jamil, T., Ly, A., Verhagen, J., Selker, R., Gronau, Q.F., Dropmann, D., Boutin, B., Meerhoff, F., Knight, P., Raj, A., Kesteren, E.-J. van, Doorn, J. van, Šmíra, M., Epskamp, S., Etz, A., Matzke, D., Jong, T. de, Bergh, D. van den Sarafoglou, A., Steingroever, H., Derks, K., Rouder, J.N., Morey, R.D., 2018. Bayesian inference for psychology. Part II: Example applications with JASP. Psychon Bull Rev 25, 58–76. https://doi.org/10.3758/s13423-017-1323-7

Weisstein, Eric W. "Bonferroni Correction." From MathWorld‐‐A Wolfram Web Resource. http://mathworld.wolfram.com/BonferroniCorrection.html

Wetzel, N., Schröger, E., Widmann, A., 2013. The dissociation between the P3a event-related potential and behavioral distraction. Psychophysiology 50, 920–930. https://doi.org/10.1111/psyp.12072

Wetzel, N., Widmann, A., Berti, S., Schröger, E., 2006. The development of involuntary and voluntary attention from childhood to adulthood: A combined behavioral and event-related potential study. Clinical Neurophysiology 117, 2191–2203. https://doi.org/10.1016/j.clinph.2006.06.717

Wetzel, N., Widmann, A., Schröger, E., 2012. Distraction and facilitation-two faces of the same coin? J Exp Psychol Hum Percept Perform 38, 664–674. https://doi.org/10.1037/a0025856

Widmann, A., Schröger, E., Wetzel, N., 2018. Emotion lies in the eye of the listener: Emotional arousal to novel sounds is reflected in the sympathetic contribution to the pupil dilation response and the P3. Biological Psychology 133, 10–17. https://doi.org/10.1016/j.biopsycho.2018.01.010

Yago, E., Escera, C., Alho, K., Giard, M.-H., Serra-Grabulosa, J.M., 2003. Spatiotemporal dynamics of the auditory novelty-P3 event-related brain potential. Cognitive Brain Research 16, 383–390. https://doi.org/10.1016/S0926-6410(03)00052-1

